# Uncertainty-aware graph representation learning with positive-unlabeled classification for biomarker discovery in peripheral artery disease

**DOI:** 10.64898/2026.05.08.723757

**Authors:** Venkat Ayyalasomayajula, Max L. Senders, Jelmer M. Wolterink, Kak Khee Yeung

## Abstract

Peripheral artery disease (PAD) is a complex vascular disorder characterized by heterogeneous molecular mechanisms and incomplete functional annotation, limiting systematic biomarker discovery. Network-based learning approaches provide a powerful framework for disease gene prioritization; however, most existing methods produce overconfident predictions without explicitly accounting for model uncertainty or structural novelty. Here, we present an uncertainty-aware framework for PAD biomarker discovery that integrates unsupervised graph representation learning, positive–unlabeled (PU) classification, ensemble prediction, and mechanistic explainability. Node embeddings were learned using multiple unsupervised graph neural network (GNN) objectives and combined with heterogeneous classifiers to generate ensemble-averaged probability estimates and epistemic uncertainty. By jointly modeling predictive confidence and embedding-space novelty, we stratified candidates into high-confidence rediscoveries and structurally novel hypotheses under explicit uncertainty control. Across eight embedding objectives and five classifiers, ensemble aggregation produced stable, well-calibrated predictions and enabled prioritization of 100 candidate PAD-associated proteins. Probability-heavy candidates clustered tightly with known PAD proteins and were enriched for established vascular and hemostatic pathways, including extracellular matrix organization, integrin signaling, coagulation, and fibrinolysis. In contrast, novelty-heavy candidates occupied distinct embedding-space regions and partitioned into multiple coherent clusters enriched for upstream regulatory and signaling processes, including G protein-coupled receptor, ephrin receptor, kinase-driven, and NF-*κ*B-associated pathways. Five-fold cross-validated comparison with established PU learning baselines demonstrated consistent improvement across all evaluation metrics (AUC 0.916 ± 0.019 vs. 0.821 ± 0.030 for the best baseline), and external validity was confirmed by significant enrichment of top candidates for related cardiovascular disease annotations (5.7× above background). Together, these results demonstrate that integrating uncertainty, novelty, and explainability enables calibrated and biologically grounded biomarker prioritization, with broad applicability to PAD and other complex diseases.

**Author summary:** Peripheral artery disease affects millions of people worldwide but remains underdiagnosed, partly because we lack reliable molecular markers to detect it early. In this study, we developed a computational framework that uses protein interaction network data to predict which proteins may be involved in PAD, even when we only know a small number of confirmed disease-associated proteins. Our approach combines graph neural network embeddings with a machine learning technique called positive-unlabeled learning, which is specifically designed for situations where you have confirmed positives but no confirmed negatives. We also quantify how confident the model is in each prediction and identify candidates that are genuinely novel compared to what is already known. Tested against established methods, our framework consistently found more known disease proteins in cross-validated evaluation. The candidates we identified map to biologically coherent pathways relevant to vascular disease, and our top predictions are enriched for proteins associated with related cardiovascular conditions, providing external validation. This work provides a principled and transparent approach to biomarker discovery that could be applied to other complex diseases with limited molecular annotations.

## Introduction

Peripheral artery disease (PAD) is a prevalent manifestation of systemic atherosclerosis that affects the lower extremities, with an estimated 236 million people affected worldwide [1]. Its prevalence is estimated to be 10 to 20% among adults over the age of 65 in community-based cohorts, and up to 30% in primary care populations over the age of 50 [2]. PAD contributes to exertional leg discomfort, functional limitation, and reduced quality of life, and is strongly associated with coronary, cerebral, and renal artery disease [3]. Individuals with PAD are at increased risk for myocardial infarction, ischemic stroke, aneurysm rupture, and vascular-related mortality, even in patients without classical symptoms [4, 5]. Although therapies such as antiplatelet agents, statins, and angiotensin-converting enzyme inhibitors have demonstrated efficacy in reducing cardiovascular events in this population [6], PAD remains underdiagnosed and undertreated [7]. Factors contributing to this gap include poor public awareness, limited physician training, the absence of reimbursement for screening, and the fact that classical intermittent claudication occurs in only a minority of patients [8]. As emphasized in major consensus statements, the under-recognition of PAD and the availability of effective therapies highlight the urgent need for improved strategies to facilitate early detection and treatment [9]. This study focuses on the identification of circulating protein biomarkers associated with PAD at the diagnostic level, encompassing the broad disease spectrum from asymptomatic ankle-brachial index abnormalities to symptomatic claudication, rather than severity staging or prediction of critical limb-threatening ischaemia (CLTI). Biomarker discovery represents a critical path toward improving PAD diagnosis and risk stratification. An ideal biomarker should demonstrate high sensitivity and specificity, correlate with disease severity, and be amenable to high-throughput assays [10, 11]. Two complementary strategies have been pursued: targeted panels of known inflammatory or metabolic markers, and unbiased mass spectrometry-based proteomic profiling [12, 13]. Despite progress, PAD biomarker discovery remains constrained by small cohorts, confounding comorbidities, technical variability, and poor reproducibility across platforms [14, 15]. These challenges motivate integrative computational frameworks capable of learning from complex, multidimensional biological data under limited supervision.

While multi-omics and machine learning methods have increasingly supported biomarker discovery, they are often limited by a scarcity of labeled data and challenges in model explainability [16, 17]. This issue is particularly acute in biological domains, where only a small subset of disease-relevant genes or proteins have been experimentally validated, while the vast majority remain unlabeled. Protein-protein interaction (PPI) networks encode the relational architecture of cellular systems and offer a principled basis for disease gene prioritization [18, 19]. In these networks, nodes represent proteins and edges capture functional or physical interactions, enabling models to exploit topological context. However, traditional supervised learning frameworks are poorly suited for this domain due to the lack of reliable negative labels.

Positive-unlabeled (PU) learning addresses this challenge by enabling classifiers to distinguish between known positives and unlabeled instances, some of which may also be positives [20, 21]. This approach is particularly well-suited for disease gene discovery, where absence of annotation does not imply irrelevance. Existing PU frameworks for disease gene prioritization, including PUDI [20], EPU [22], ProDiGe [23], and NIAPU [24], have demonstrated the value of this paradigm but share important limitations: none explicitly quantify model uncertainty, and none provide mechanisms for distinguishing high-confidence rediscoveries from structurally novel candidates.

Here, we address these limitations with a three-phase framework that integrates unsupervised graph representation learning, uncertainty-aware PU classification, and post-hoc mechanistic explainability for PAD biomarker discovery. Our contributions are: (i) a modular uncertainty-aware PU learning framework for biomarker discovery in weakly labeled molecular networks; (ii) a dual selection strategy combining confidence-driven and novelty-driven candidate prioritization; (iii) group-level consensus explainability using GNNExplainer; and (iv) systematic five-fold cross-validated comparison with established PU learning baselines with external validity assessment.

## Methods

### PPI network and protein representations

We constructed a multimodal PPI network focused on PAD using high-confidence interactions from STRING v12.0 [25], retaining only human interactions with a combined confidence score above 0.7 and extracting the largest connected component, yielding approximately 7,600 protein nodes. PAD-associated proteins were identified through rigorous literature review of peer-reviewed case-control plasma proteomics studies comparing PAD patients across disease severity stages to healthy controls, yielding 68 proteins with experimentally supported PAD associations, labelled as positives. All remaining proteins were treated as unlabelled, consistent with the PU learning setting. An overview of the complete framework is provided in Fig 1.

**Fig 1.**
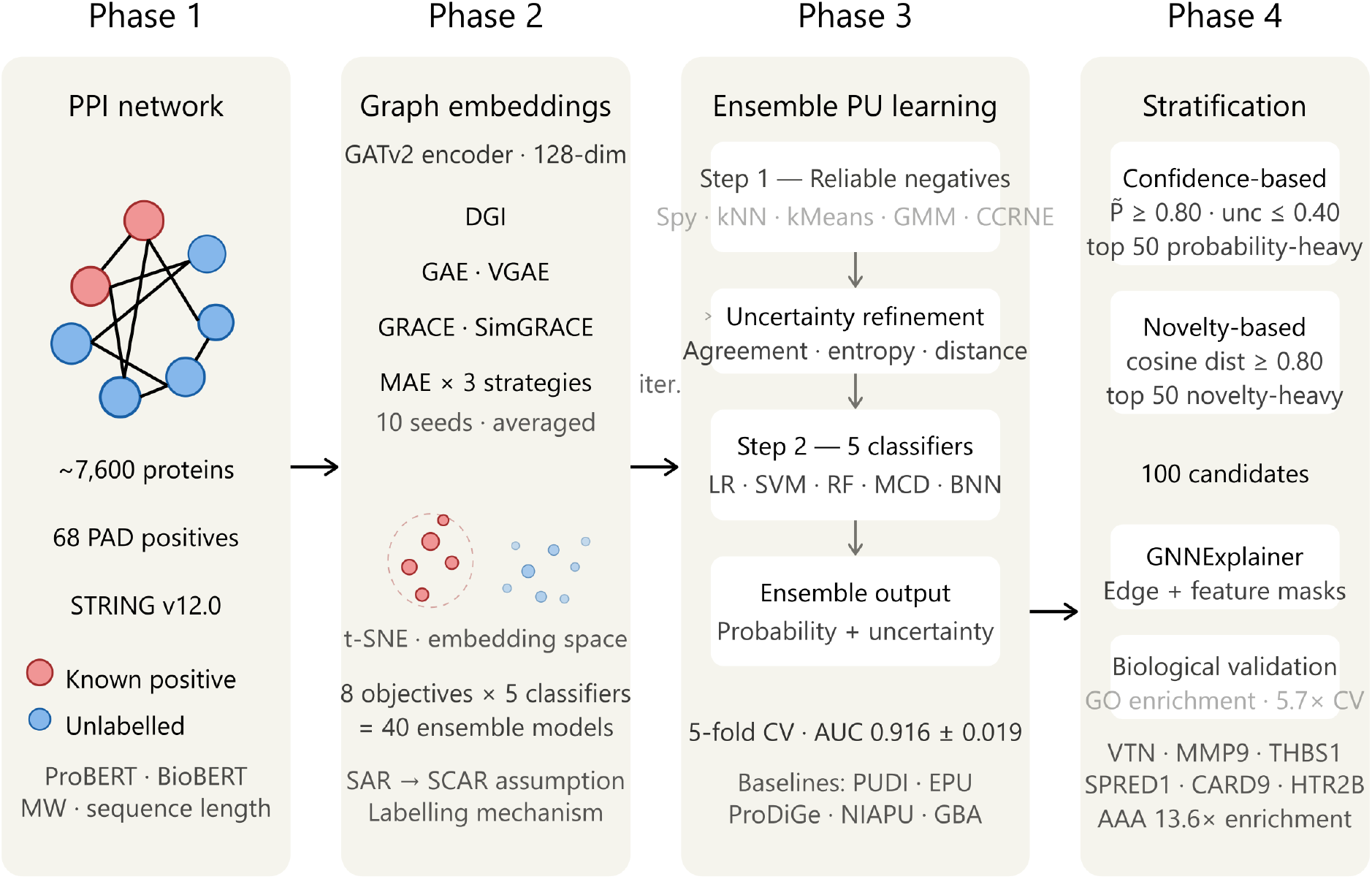
Uncertainty-aware framework for PAD biomarker discovery. The pipeline comprises four phases. **(1) PPI network:** A protein–protein interaction (PPI) network is constructed from STRING v12.0 (7,600 nodes). Known PAD-positive proteins (*n* = 68, red) are identified from plasma proteomics literature; remaining proteins are unlabelled (blue). Node feature vectors combine ProBERT sequence embeddings, BioBERT functional annotation embeddings, molecular weight, and sequence length. **(2) Graph embeddings:** A shared GATv2 encoder produces 128-dimensional embeddings under eight unsupervised objectives: DGI, GAE, VGAE, GRACE, SimGRACE, and MAE with three masking strategies. Each objective is trained with 10 random seeds and averaged. Labelling is assumed SAR, reduced to SCAR following Bekker & Davis (2020). **(3) Ensemble PU learning:** Reliable negatives are identified via five heuristics (Spy, *k*NN, *k*-means, GMM, CCRNE) and refined by uncertainty scoring. Five classifiers - logistic regression (LR), support vector machine (SVM), random forest (RF), Monte Carlo Dropout (MCD), and Bayesian neural network (BNN), are trained iteratively across 40 ensemble instances, achieving AUC 0.916 ± 0.019 in 5-fold cross-validation versus 0.821 ± 0.030 for the best baseline. **(4) Candidate stratification:** 100 novel candidates are selected by confidence (top 50, 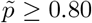) and cosine novelty (top 50, dist≥ 0.80). Post-hoc explainability uses GNNExplainer; biological validation shows 5.7× cardiovascular enrichment (AAA 13.6×). PAD: peripheral artery disease; GATv2: Graph Attention Network v2; DGI: Deep Graph Infomax; GAE/VGAE: Graph Autoencoder/Variational GAE; GRACE/SimGRACE: graph contrastive learning methods; MAE: Masked Autoencoder; SAR/SCAR: Selected At/Completely At Random; LR: logistic regression; SVM: support vector machine; RF: random forest; MCD: Monte Carlo Dropout; BNN: Bayesian neural network; GMM: Gaussian mixture model; CCRNE: cluster-complementary reliable negative extraction; AAA: abdominal aortic aneurysm.

Each protein node was encoded using a multimodal feature representation combining ProBERT sequence embeddings [26] and BioBERT functional annotation embeddings [27], concatenated with scalar biochemical attributes including molecular weight and sequence length.

### Labelling mechanism and PU learning assumptions

A critical consideration in PU learning is the mechanism by which positive examples are selected for labelling [21]. Our positive set was derived from experimental plasma proteomics studies, introducing systematic labelling bias: proteins are more likely to be identified and reported if they have prior biological plausibility, are detectable at sufficient plasma abundance, or belong to well-studied pathways. This corresponds to a Selected At Random (SAR) labelling mechanism, where the probability of a protein being labelled depends on its observed attributes.

However, following [21], we argue that SAR can be reduced to Selected Completely At Random (SCAR) in our setting, because the attributes driving labelling, experimental detectability, prior literature coverage, and pathway familiarity are orthogonal to the attributes used for classification, namely unsupervised GNN-derived topological embeddings from the PPI network structure. A protein’s position in the PPI network topology is not what determines whether it is studied in a plasma proteomics experiment. Under this reduction, recall-based cross-validated evaluation is valid, and our evaluation metrics provide unbiased estimates of method performance.

We further assume a single-training-set scenario [21], whereby both positive and unlabeled proteins originate from the same PPI network dataset. This is the most common scenario in PU learning applications and is satisfied here since all nodes, regardless of label status, are drawn from the same STRING-derived network.

We additionally assume separability, that a classifier exists capable of distinguishing PAD-associated from non-associated proteins in embedding space and smoothness, that proteins with similar embedding representations have similar probabilities of PAD association. The structured separation of known PAD proteins from unlabeled background nodes in t-SNE projections of the learned embeddings (S1 Fig) provides empirical support for both assumptions.

### Unsupervised graph representation learning

We trained a two-layer Graph Attention Network v2 (GATv2) [28] as the shared encoder across all unsupervised objectives, producing 128-dimensional node embeddings. GATv2 improves over standard GAT [29] by applying the attention mechanism after the linear transformation, enabling more expressive neighbour weighting. Full architectural details and training objectives are provided in the Supplementary Methods (S1 Text).

We evaluated eight self-supervised objectives applied to the same backbone: Deep Graph Infomax (DGI) [30], Graph Autoencoder (GAE) and Variational GAE (VGAE) [31], GRACE [32], SimGRACE [33], and Masked Autoencoder (MAE) with three masking strategies (random, degree-based, adaptive) [34]. Each model was run for 10 independent initialisations, with final embeddings obtained by element-wise averaging across runs.

### Positive-unlabeled learning framework

We adopted a two-step PU learning strategy [21]. Step 1 identifies reliable negatives (RNs) from the unlabelled pool using an ensemble of five complementary heuristics: the Spy method [**?**], *k*-means distance filter, *k*-NN distance ranking, Gaussian mixture model scoring, and cluster-complementary RN extraction (CCRNE). RN candidates are subsequently refined by joint scoring of heuristic agreement, entropy-based uncertainty, and distance to known positives. Step 2 trains five classifiers iteratively on labelled positives and refined RNs: logistic regression, SVM, random forest, MC Dropout [35], and Bayesian neural networks [36]. The iterative loop expands the negative set with high-confidence unlabelled negatives across a maximum of 10 iterations. Full algorithmic details and all equations are provided in S1 Text.

### Ensemble prediction and uncertainty quantification

Final ensemble scores were obtained by averaging predictions across all eight embedding objectives and five classifiers (40 model instances total), yielding stable, well-calibrated probability and uncertainty estimates for each protein node. For MC Dropout and Bayesian neural network classifiers, 30 stochastic forward passes were drawn to obtain mean predicted probabilities and epistemic uncertainty via predictive entropy. For deterministic classifiers, probability outputs were used directly with scalar entropy estimation. Full equations are provided in S1 Text.

### Candidate selection strategy

Following ensemble prediction, candidates were prioritised using two complementary strategies. **Confidence-based selection** (probability-heavy) retained nodes with ensemble mean probability 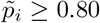 and epistemic uncertainty ≤ 0.40, selecting the top 50 by descending probability. **Novelty-based selection** (cosine-heavy) quantified structural dissimilarity from known positives via mean cosine distance in embedding space, 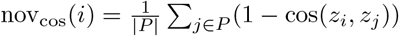, retaining nodes with 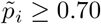, uncertainty ≤ 0.40, and nov_cos_(*i*) ≥ 0.80, and selecting the top 50 by descending novelty. The final candidate set comprised 100 novel proteins spanning both selection regimes.

### Mechanistic explainability

We employed GNNExplainer [37] to generate post-hoc explanations for all prioritised candidates. For each candidate node, GNNExplainer learns edge and feature masks by maximising mutual information between the original prediction and the masked inputs, yielding a minimal explanatory subgraph. Explanations were generated across all model instances and aggregated by averaging, producing consensus explanations. Explanation stability was quantified as the variance of feature-importance masks across model instances; explanation faithfulness was assessed by the concentration of importance in top-ranked features [38]. Group-level consensus subnetworks were constructed by retaining the top max(100, 1%) edges ranked by mean explanatory importance across all candidates within each group. Feature versus topology importance was quantified by comparing aggregate node-level feature mask mass against edge mask mass, classifying predictions as feature-driven, topology-driven, or balanced. Full configuration details in S1 Text.

### Baseline methods and evaluation

We compared our framework against five established methods: PUDI [20] (spy method + single LR classifier), EPU [22] (ensemble PU without uncertainty), ProDiGe [23] (bagging-based PU), NIAPU [24] (network-informed adaptive PU), and GBA (guilt-by-association via cosine similarity to known positives [39]). All methods operated on the same ensemble-averaged embeddings as input, ensuring a fair comparison of learning strategies given identical representations.

Evaluation followed the cross-validated recall framework of [21], using 5-fold cross-validation on the 68 known PAD positives. In each fold, approximately 20% of positives were held out as a test set; all methods were trained on the remaining 80% positives plus the unlabelled pool. Performance was assessed using AUC-ROC, average precision (AP), Precision@*K*, and Recall@*K* for *K* ∈ {25, 50, 100, 200}. External validity was assessed by computing the enrichment of top-100 candidates for proteins annotated to related cardiovascular diseases (AAA, AMI, Stroke, Heart Failure) in our dataset, relative to the background annotation rate across all unlabelled nodes.

## Results

### Embedding quality and reliable negative selection

Across eight unsupervised objectives, all methods produced 128-dimensional embeddings with non-degenerate variance structure. The effective dimensionality required to explain 90% of variance ranged from 4 to 99 dimensions across methods, indicating complementary representational properties. Embedding stability was consistently low (mean node-wise standard deviation 0.048–0.074), confirming reproducible representations despite stochastic training (S1 Fig). This structured separation provides empirical support for the separability and smoothness assumptions underlying our PU learning framework.

Reliable negative selection yielded pools with substantial cross-heuristic agreement: 55–70% of candidates were supported by at least two independent heuristics, while fewer than 20% were supported by a single method. Iterative PU training converged within 5–10 iterations, expanding reliable negative sets to 1,360–2,166 nodes depending on classifier and embedding method.

### Ensemble predictions and candidate stratification

Ensemble aggregation yielded well-calibrated probability and uncertainty estimates across the protein network (Fig 2A). The ensemble distribution was strongly right-skewed, with a distinct high-confidence subset exhibiting both high predicted probability and low epistemic uncertainty, enabling clear separation of predicted positives from the background.

**Fig 2.**
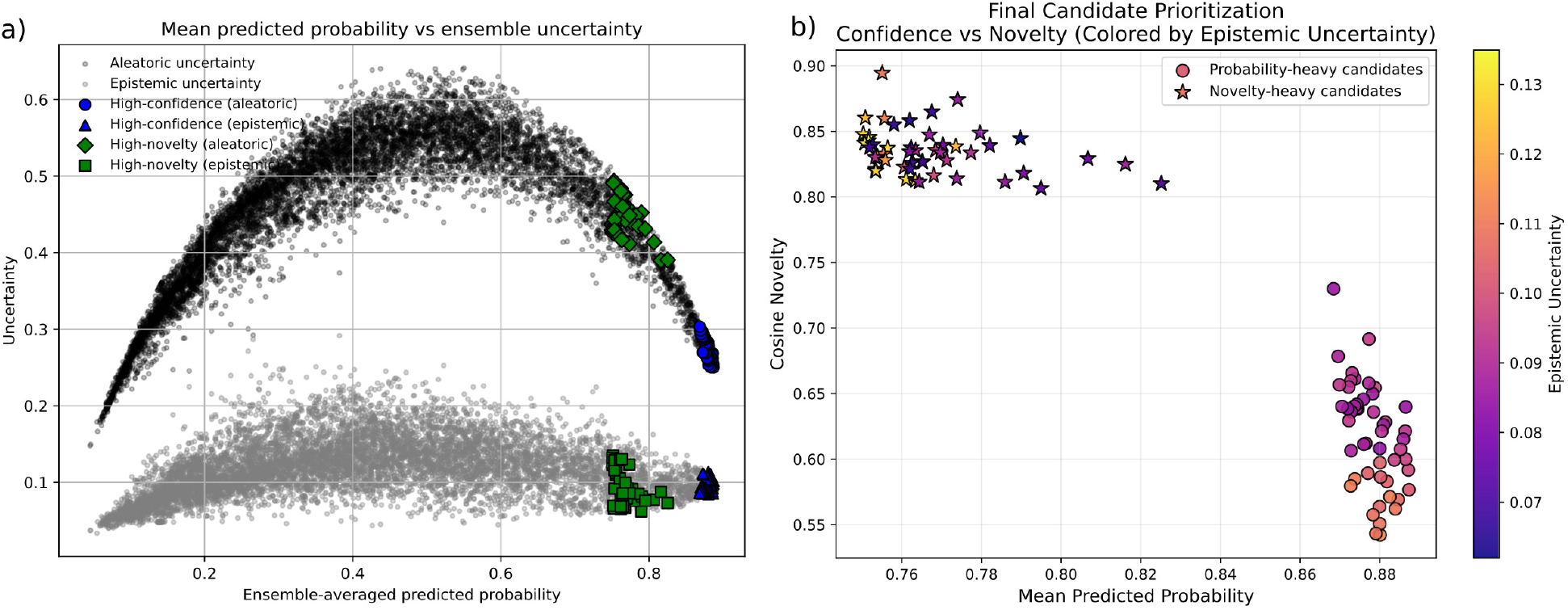
Ensemble prediction confidence, uncertainty, and candidate stratification. (A) Ensemble-averaged predicted probability versus epistemic uncertainty for all nodes. Gray points denote background nodes; highlighted points indicate final prioritized candidates separated into probability-heavy and cosine-novelty-heavy sets. (B) Confidence–novelty landscape for prioritized candidates, showing ensemble mean probability versus cosine novelty. Probability-heavy candidates (circles) and novelty-heavy candidates (stars) are colored by ensemble epistemic uncertainty.

Joint consideration of ensemble confidence and embedding-space novelty revealed two complementary candidate regimes (Fig 2B). Probability-heavy candidates concentrated in regions of high ensemble probability and low uncertainty, reflecting strong classifier support and close proximity to known PAD proteins. Cosine-heavy candidates displayed substantially greater embedding-space separation from known PAD proteins while retaining moderate-to-high predicted probability, indicating structural novelty without loss of predictive support. The final prioritised set comprised 100 novel candidates, evenly split between both groups, as summarised in Table 1.

**Table 1.** Selected prioritised novel PAD biomarker candidates. Top 10 probability-heavy (P) and top 5 cosine-heavy (C) candidates with ensemble confidence, epistemic uncertainty, cosine novelty, and representative biological pathway. Full list of all 100 candidates in S1 Table.

| Rank | Gene | Group | Mean P | Uncertainty | Cosine Nov. | Key Pathway |
| --- | --- | --- | --- | --- | --- | --- |
| 1 | VTN | P | 0.887 | 0.102 | 0.577 | ECM / Integrin |
| 2 | ITGAV | P | 0.887 | 0.098 | 0.592 | Integrin signalling |
| 3 | HABP2 | P | 0.886 | 0.085 | 0.640 | Coagulation |
| 4 | ITGB6 | P | 0.886 | 0.097 | 0.600 | Integrin signalling |
| 5 | ITGA2 | P | 0.886 | 0.092 | 0.621 | Cell-matrix adhesion |
| 6 | THBS1 | P | 0.883 | 0.111 | 0.571 | ECM organisation |
| 7 | MMP9 | P | 0.879 | 0.108 | 0.543 | ECM remodelling |
| 8 | VWF | P | 0.873 | 0.111 | 0.580 | Haemostasis |
| 9 | APOH | P | 0.870 | 0.089 | 0.678 | Coagulation |
| 10 | JAK3 | P | 0.868 | 0.086 | 0.730 | JAK-STAT signalling |
| 51 | SPRED1 | C | 0.825 | 0.073 | 0.810 | MAPK regulation |
| 52 | PRKCD | C | 0.790 | 0.062 | 0.845 | Kinase signalling |
| 53 | MAP3K7 | C | 0.782 | 0.076 | 0.839 | NF-κB / MAPK |
| 54 | INPP5D | C | 0.774 | 0.084 | 0.874 | PI3K signalling |
| 55 | HTR2B | C | 0.755 | 0.110 | 0.894 | GPCR signalling |

### Baseline comparison

Five-fold cross-validated evaluation demonstrated consistent superiority of our framework over all baselines across all metrics (Table 2). Our method achieved a mean AUC of 0.916 ± 0.019, compared to 0.821 ± 0.030 for the best-performing baseline (GBA), representing a margin of approximately 10 AUC points. Average precision was 0.032 ± 0.013 versus 0.019 ± 0.007 for GBA. Recall@100 reached 0.247 ± 0.081, substantially exceeding GBA (0.189 ± 0.083) and all PU learning baselines (range 0.103–0.133). The consistent improvement across all *K* values and both precision and recall metrics confirms that the uncertainty-aware ensemble framework provides more reliable candidate prioritisation than existing approaches given identical input representations.

**Table 2.** Five-fold cross-validated comparison with baseline methods. All methods operate on identical ensemble-averaged GNN embeddings. Values are mean ± standard deviation across 5 folds. Bold indicates best performance.

| Method | AUC | AP | P@50 | R@50 | P@100 | R@100 |
| --- | --- | --- | --- | --- | --- | --- |
| PUDI [20] | 0.798±0.046 | 0.016±0.008 | 0.016±0.008 | 0.059±0.030 | 0.014±0.010 | 0.103±0.074 |
| EPU [22] | 0.813±0.054 | 0.014±0.006 | 0.012±0.016 | 0.044±0.058 | 0.016±0.010 | 0.118±0.073 |
| ProDiGe [23] | 0.811±0.053 | 0.016±0.008 | 0.012±0.016 | 0.044±0.058 | 0.018±0.012 | 0.133±0.086 |
| NIAPU [24] | 0.800±0.049 | 0.014±0.004 | 0.012±0.010 | 0.044±0.036 | 0.018±0.013 | 0.131±0.095 |
| GBA [39] | 0.821±0.030 | 0.019±0.007 | 0.032±0.016 | 0.117±0.058 | 0.026±0.012 | 0.189±0.083 |
| **Ours** | **0.916±0.019** | **0.032±0.013** | **0.048±0.027** | **0.175±0.095** | **0.034±0.012** | **0.247±0.081** |

### External validity

Among the top 100 novel candidates, 29% carried annotations for at least one related cardiovascular disease (AAA, AMI, Stroke, or Heart Failure), compared to a background rate of 5.1% across all unlabelled nodes, a 5.7-fold enrichment. Enrichment was most pronounced for AAA (13.6×), consistent with shared atherosclerotic mechanisms between PAD and AAA, followed by AMI (3.2×), Stroke (1.8×), and Heart Failure (1.7×). This pattern validates that our framework preferentially identifies proteins with broader cardiovascular relevance rather than arbitrary high-scoring nodes.

### Functional enrichment

Functional enrichment analysis revealed distinct but biologically coherent signatures for the two candidate groups (Fig 3). Probability-heavy candidates were significantly enriched for adhesion-and haemostasis-related processes, including cell adhesion mediated by integrins, integrin-mediated signalling, regulation of blood coagulation, and fibrinolysis, closely overlapping with pathways enriched among known PAD proteins. Novelty-heavy candidates were enriched for intracellular signalling and regulatory processes, including NF-*κ*B signalling, ephrin receptor signalling, peptidyl-tyrosine phosphorylation, MAPK cascade regulation, and protein linear polyubiquitination, consistent with upstream regulatory control and immune-vascular signalling.

**Fig 3.**
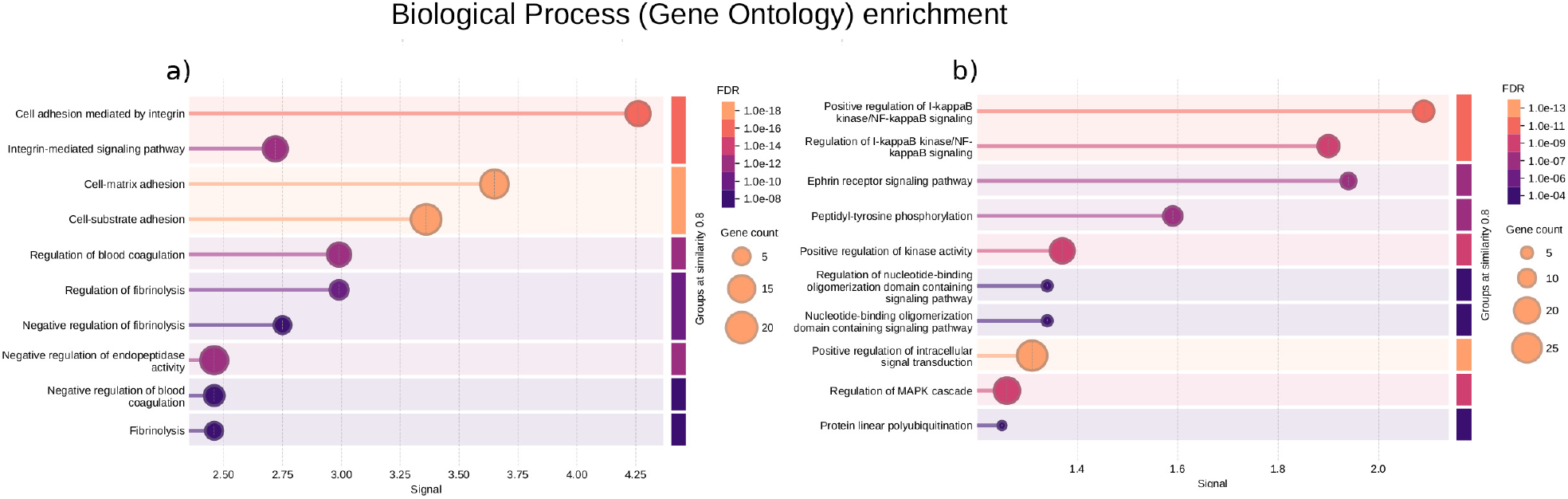
Functional enrichment of prioritized candidate biomarkers. Gene Ontology biological process enrichment analysis for probability-heavy (left) and novelty-heavy (right) candidate sets. Probability-heavy candidates are enriched for adhesion-, coagulation-, and extracellular matrix-related processes, whereas novelty-heavy candidates are enriched for intracellular signaling and regulatory pathways including NF-*κ*B, MAPK, and ephrin receptor signaling. Only the top enriched terms for each group are shown.

### Clustering analysis

Clustering revealed marked structural differences across candidate groups. Known PAD proteins exhibited diffuse but coherent organisation, with 27 of 68 assigned to two clusters dominated by vascular remodelling and haemostatic processes. Probability-heavy candidates showed markedly consolidated structure, with 44 of 50 assigned to two clusters mirroring known PAD organisation. In contrast, novelty-heavy candidates exhibited plural organisation across four medium-sized clusters corresponding to G protein-coupled receptor signalling, ephrin receptor signalling, kinase-driven transduction, and innate immune pathways. This multi-cluster structure confirms that novelty-based predictions capture mechanistically diverse hypotheses distributed across multiple independent regulatory neighbourhoods.

### Mechanistic explainability

Explanation stability was consistently low for both candidate groups (feature mask variance range 0.03–0.07), confirming reproducible explanatory behaviour. PAD neighbourhood context differed markedly: probability-heavy candidates showed broader distributions of PAD-labelled neighbours, while novelty-heavy candidates were predominantly characterised by zero or near-zero PAD-labelled neighbours (Fig 4).

**Fig 4.**
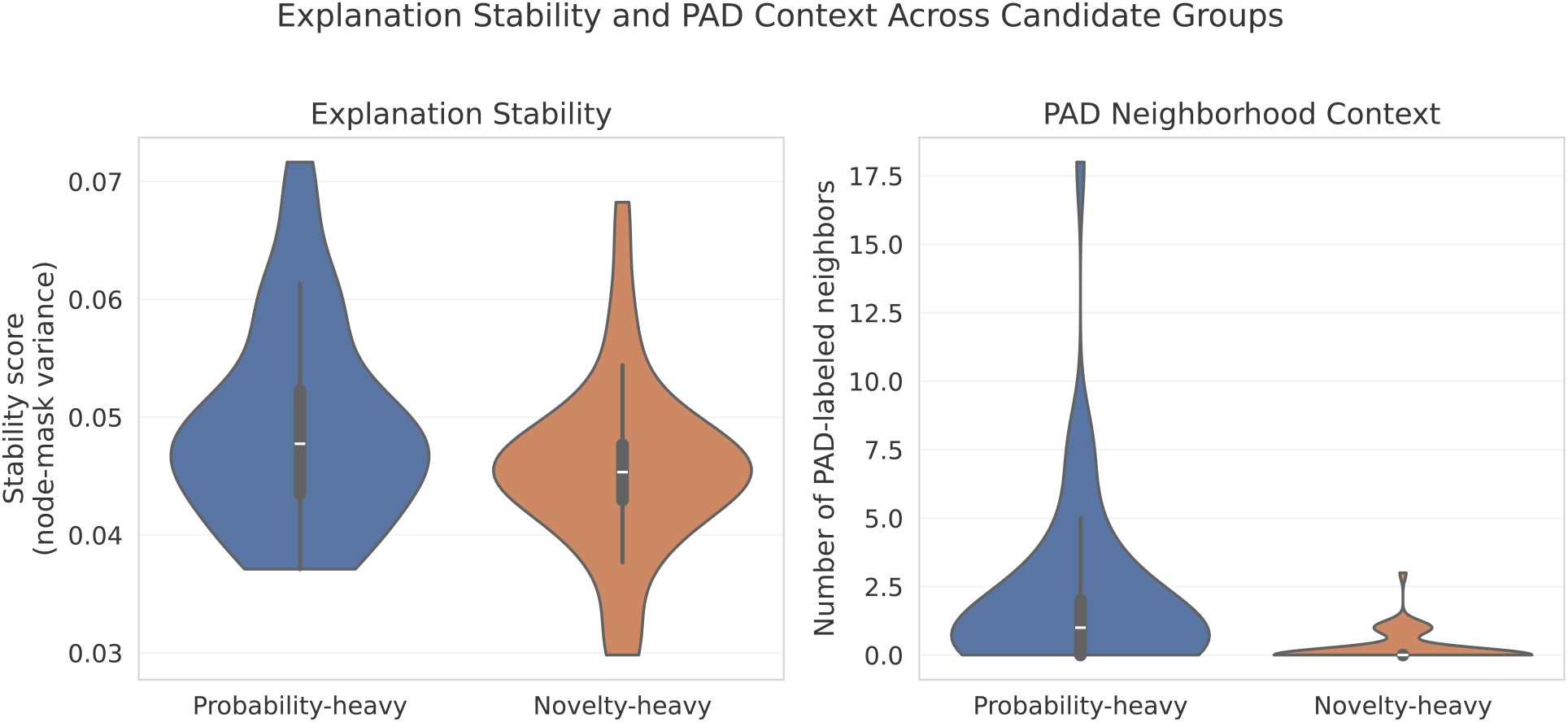
Explanation stability and PAD neighborhood context across candidate groups. (A) Distribution of explanation stability scores, quantified as the variance of node-level feature masks across repeated GNNExplainer runs. Lower values indicate more stable explanations. (B) Distribution of PAD neighborhood support, measured as the number of directly connected PAD-labeled neighbors in the protein interaction network.

Group-level consensus subnetworks highlighted recurrent interaction patterns. The probability-heavy consensus network contained 116 nodes including 11 candidates and 8 known PAD proteins, while the novelty-heavy consensus network comprised 170 nodes including 44 candidates but only 3 known PAD proteins (Fig 5), confirming that probability-heavy predictions are anchored in established PAD neighbourhoods while novelty-heavy predictions emerge from distinct network regions. Feature versus topology analysis revealed that topology dominates predictions for both groups, with topology-driven predictions exceeding 60% in both groups, underscoring that PPI network structure rather than individual protein features is the primary driver of confident prioritisation.

**Fig 5.**
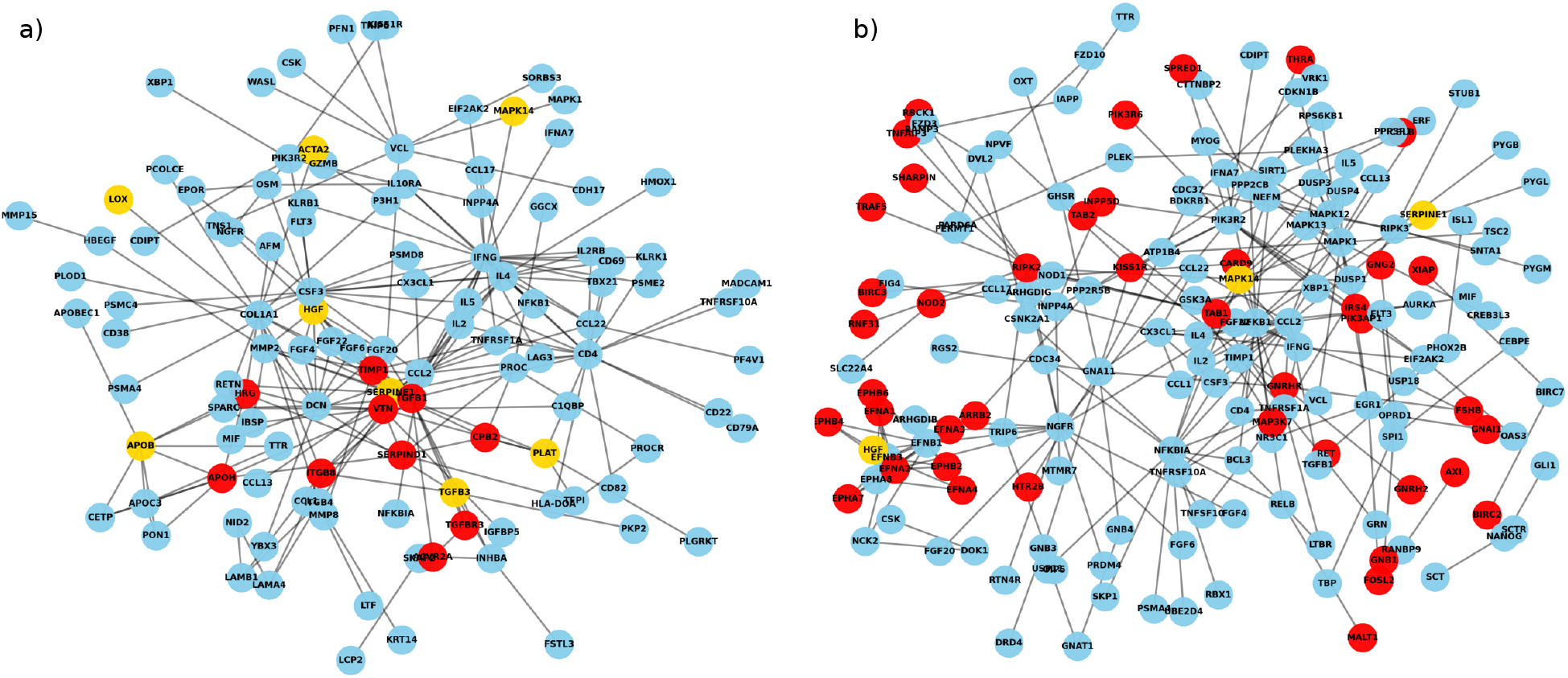
Group-level consensus interaction networks derived from aggregated GNNExplainer edge attributions. Left: probability-heavy candidates; right: novelty-heavy candidates. Red nodes denote prioritized candidates, yellow nodes indicate known PAD-associated proteins, and gray nodes represent other proteins. Only candidates participating in shared high-importance interactions appear in the consensus graphs.

Feature group importance analysis revealed that no single feature modality dominated predictions; instead, importance was distributed across all six feature groups, with molecular weight and sequence length features showing slightly higher relative importance in probability-heavy candidates, while novelty-heavy candidates exhibited a more uniform distribution across all groups (S3 Fig). This confirms that predictions are not driven by any single protein property but reflect a holistic integration of multimodal features within the network context.

## Discussion

Here is the complete rewritten discussion with all additions integrated, keeping your existing text intact: latex

### Uncertainty-aware prioritisation and baseline comparison

Our framework advances network-based disease gene prioritisation by moving beyond binary classification toward calibrated, risk-aware hypothesis generation. The cross-validated comparison against PUDI, EPU, ProDiGe, NIAPU, and GBA demonstrates that explicit uncertainty quantification and ensemble aggregation consistently improve recovery of known disease-associated proteins, with a margin of approximately 10 AUC points over the best-performing baseline. Most existing frameworks implicitly treat all high-scoring candidates as equally reliable despite substantial uncertainty arising from model choice, training stochasticity, and incomplete biological knowledge [31, 40]. By integrating ensemble-averaged predictive confidence with epistemic uncertainty [36], our approach enables principled separation of well-supported predictions from uncertain extrapolations, treating uncertainty as an informative signal that contextualises each candidate’s reliability [41]. The dual selection strategy further enriches this framework by introducing novelty as a complementary axis: probability-heavy candidates represent high-confidence rediscoveries aligned with established PAD biology, while novelty-heavy candidates occupy structurally distinct embedding regions yet retain moderate predictive support. This calibrated prioritisation is essential for translational settings, where the cost of false confidence often outweighs that of cautious exploration [42, 43].

### Biological interpretation of candidate groups

Rather than revealing a single dominant explanatory pattern, our analyses exposed substantial mechanistic heterogeneity across candidates. Some probability-heavy candidates, such as VTN, were embedded within explanatory subnetworks densely connected to known PAD-associated proteins. Others, including LIFR, exhibited explanatory subgraphs with limited direct overlap with known PAD proteins, yet highlighted coherent interaction neighbourhoods, indicating that proximity to known disease genes is not a prerequisite for confident prediction [39]. A similar spectrum was observed among novelty-heavy candidates, with some such as RBCK1 displaying connections to other candidates and indirect PAD links, while others including HTR2B were supported primarily by distinct local structures. Topology dominates predictions across both groups, underscoring that network context rather than individual protein properties drives confident prioritisation.

Probability-heavy candidates recapitulate well-characterised PAD pathways, extracellular matrix organisation, integrin-mediated adhesion, coagulation, and fibrinolysis [2, 44], and consolidate around two dominant clusters mirroring known PAD organisation. Novelty-heavy candidates highlight complementary regulatory processes, G protein-coupled receptor signalling, ephrin receptor signalling, NF-*κ*B pathways, and MAPK cascades [45–47], distributed across multiple coherent mechanistic modules, consistent with upstream regulatory control and immune-vascular crosstalk [48]. The hierarchical organisation of candidates revealed by clustering analysis provides structural corroboration for these functional distinctions. Probability-heavy candidates consolidated around two dominant clusters mirroring the organisation of known PAD-associated proteins, indicating convergence onto a limited number of shared disease-supporting mechanisms consistent with their interpretation as high-confidence rediscoveries. In contrast, novelty-heavy candidates fragmented across four independent mechanistic clusters corresponding to G protein-coupled receptor signalling, ephrin receptor signalling, kinase-driven transduction, and innate immune pathways. This plural organisation confirms that novelty-based prioritisation does not identify a single alternative disease pathway but instead captures multiple parallel regulatory layers that may modulate vascular inflammation and immune-vascular crosstalk in PAD [48]. The structural contrast between consolidated and fragmented clustering thus provides independent network-level evidence that the confidence-novelty axis separates candidates along a biologically meaningful dimension.

Among individual probability-heavy candidates, several warrant specific biological note. Vitronectin (VTN), the top-ranked candidate, is an extracellular matrix glycoprotein that mediates cell-matrix interactions through integrin binding and plays a direct role in platelet aggregation and coagulation cascade regulation [13], processes centrally dysregulated in PAD. Matrix metalloproteinase 9 (MMP9) is a well-established driver of atherosclerotic plaque instability through extracellular matrix degradation and fibrous cap thinning, with elevated plasma levels documented in vulnerable plaques [49]. Thrombospondin-1 (THBS1), a multifunctional matricellular glycoprotein, regulates platelet aggregation, vascular remodelling, and TGF-*β* activation, and has been shown to be upregulated in aortic aneurysm tissue, consistent with our external validity finding of strong AAA enrichment among top candidates [50]. Together, these examples demonstrate that probability-heavy candidates are not generic high-scoring nodes but proteins with mechanistically coherent roles in vascular pathophysiology.

### External validity and network-level evidence

The biological plausibility of prioritised candidates was independently supported by cross-disease enrichment analysis. Among the top 100 novel candidates, 29% carried annotations for at least one related cardiovascular disease (AAA, AMI, Stroke, or Heart Failure), compared to a background rate of 5.1% across all unlabelled nodes, a 5.7-fold enrichment. Enrichment fold-change was computed as the ratio of the annotation rate among top-100 candidates to the background annotation rate across all unlabelled nodes. Enrichment was most pronounced for AAA (13.6×), which is biologically coherent: PAD and AAA share underlying atherosclerotic pathophysiology, overlapping haemostatic dysfunction, and common inflammatory mediators, yet their molecular characterisation has largely proceeded independently [51, 52]. The strong enrichment of our top candidates for AAA-associated proteins therefore suggests that our framework recovers proteins operating at the intersection of these related vascular conditions, potentially identifying shared mechanistic nodes that could serve as targets across the atherosclerotic disease spectrum. The more modest but consistent enrichment for AMI (3.2×), Stroke (1.8×), and Heart Failure (1.7×) further confirms that top candidates are not arbitrary high-scoring nodes but proteins with broader cardiovascular relevance.

The finding that topology dominates predictions across both candidate groups has important implications for network-based biomarker discovery more broadly. Across all 100 candidates, topology-driven predictions exceeded 60% in both probability-heavy and novelty-heavy groups, indicating that a protein’s structural role in the interaction network, its connectivity patterns, neighbourhood composition, and topological position, is more predictive of disease association than its individual molecular properties. This aligns with and quantitatively extends the guilt-by-association principle underlying network medicine [39], providing evidence that topological context consistently outweighs feature-level signals even within an uncertainty-aware ensemble framework. For future biomarker discovery efforts, this finding argues strongly for graph-based representations over feature-only approaches, and suggests that network topology should be treated as a primary rather than auxiliary signal in disease gene prioritisation tasks.

### Translational implications

Beyond methodological contributions, our framework has practical implications for vascular biology research. The dual candidate output, high-confidence rediscoveries and structurally novel hypotheses, is designed to support two distinct downstream workflows. Probability-heavy candidates, enriched for established vascular pathways and closely embedded within known PAD interaction neighbourhoods, represent the most tractable targets for experimental validation through targeted proteomics or functional assays in existing PAD cohorts. Novelty-heavy candidates, distributed across independent regulatory modules and enriched for upstream signalling processes, are better suited for hypothesis-generating studies such as pathway perturbation screens or multi-omics integration in new cohorts. The uncertainty estimates accompanying each candidate provide an explicit prioritisation signal for resource allocation: candidates with high confidence and low epistemic uncertainty warrant immediate experimental follow-up, while those with higher uncertainty may benefit from additional computational validation before committing experimental resources. More broadly, the framework is disease-agnostic and directly applicable to any complex disease characterised by sparse molecular annotations and a well-curated PPI network, including abdominal aortic aneurysm, coronary artery disease, and heart failure, conditions for which our external validity results suggest partial mechanistic overlap with the identified candidates.

### Limitations and future directions

Several limitations should be acknowledged. The framework relies on a static global PPI network that is incomplete and biased toward well-studied proteins; incorporation of tissue-specific or context-dependent interaction networks would improve biological fidelity. While ensemble learning stabilises predictions and quantifies epistemic uncertainty, it does not capture all sources of biological variability such as disease stage or patient heterogeneity; integrating multi-omics data or longitudinal cohorts could further refine uncertainty estimates. Candidate prioritisation remains hypothesis-generating rather than causal, and experimental confirmation is required to establish direct involvement of prioritised candidates in PAD pathogenesis. Finally, GNNExplainer was applied to unsupervised graph encoders rather than directly to PU classifiers, providing an indirect but consistent view of prediction drivers; future work could explore classifier-specific or counterfactual explainability methods to more directly interrogate decision boundaries.

## Conclusion

We introduced an uncertainty-aware, graph-based framework for biomarker discovery in PAD that integrates unsupervised representation learning, ensemble PU classification, and mechanistic explainability. By jointly modeling predictive confidence, epistemic uncertainty, and embedding-space novelty, the framework enables calibrated hypothesis generation under realistic uncertainty. Cross-validated comparison with established PU learning baselines confirmed consistent improvement across all evaluation metrics, and external validity assessment demonstrated significant enrichment of top candidates for related cardiovascular disease annotations. Explanation-based analyses revealed that topology drives predictions and that confident and exploratory candidates arise from structurally distinct network regions, highlighting multiple mechanistic routes through which PAD-associated signals emerge. The proposed framework provides a generalisable and risk-aware strategy for prioritising candidates in diseases characterised by heterogeneous mechanisms and incomplete annotations, with broad applicability beyond PAD to other complex diseases.

## Supporting information

**S1 Text. Supplementary Methods**. Full GATv2 architectural details and training objectives for all eight unsupervised embedding methods; hyperparameter configurations; detailed PU learning algorithm including RN identification equations, uncertainty refinement procedure, and iterative training loop with all thresholds; GNNExplainer configuration; stability and faithfulness metric definitions; full hyperparameter table.

**S1 Fig. Embedding stability and t-SNE projection**. (A) Cross-run embedding stability across all eight unsupervised methods, quantified as mean node-wise standard deviation across random initialisations. (B) Representative two-dimensional t-SNE projection of GAE node embeddings with candidate stratification. Known PAD-associated proteins are shown in red, probability-heavy candidates in blue, and novelty-heavy candidates in green.

**S2 Fig. Representative GNNExplainer subgraphs**. Shown are two probability-heavy candidates (VTN, LIFR) and two novelty-heavy candidates (HTR2B, RBCK1). Node colors indicate candidate proteins (orange), other candidates (yellow), known PAD-associated proteins (red), and background proteins (gray).

**S3 Fig. Feature vs. topology importance**. Full feature versus topology dominance analysis for all 100 candidates under both elbow-based and density-based importance methods.

**S1 Table. Full candidate list**. Complete ranked list of all 100 prioritised novel PAD biomarker candidates with ensemble probability, epistemic uncertainty, cosine novelty, support count, and group assignment.

**S2 Table. Pathway enrichment results**. Complete Gene Ontology biological process enrichment tables for probability-heavy and novelty-heavy candidate groups.

**S3 Table. Clustering analysis**. Full cluster assignments and pathway annotations for known PAD proteins, probability-heavy candidates, and novelty-heavy candidates.

## Acknowledgments

We thank Lotte Rijken, Victor Janssen, and Patryk Rygiel for helpful and intriguing discussions.

## Author contributions

Conceptualization: VA, KKY. Formal analysis: VA. Investigation: VA. Methodology: VA, JMW. Project administration: KKY. Software: VA. Supervision: JMW, KKY. Visualization: VA. Writing – original draft: VA. Writing – review & editing: VA, MS, JMW, KKY.

## Supplementary Material

Uncertainty-aware graph representation learning with positive-unlabeled classification for biomarker discovery in peripheral artery disease

**Fig S1.**
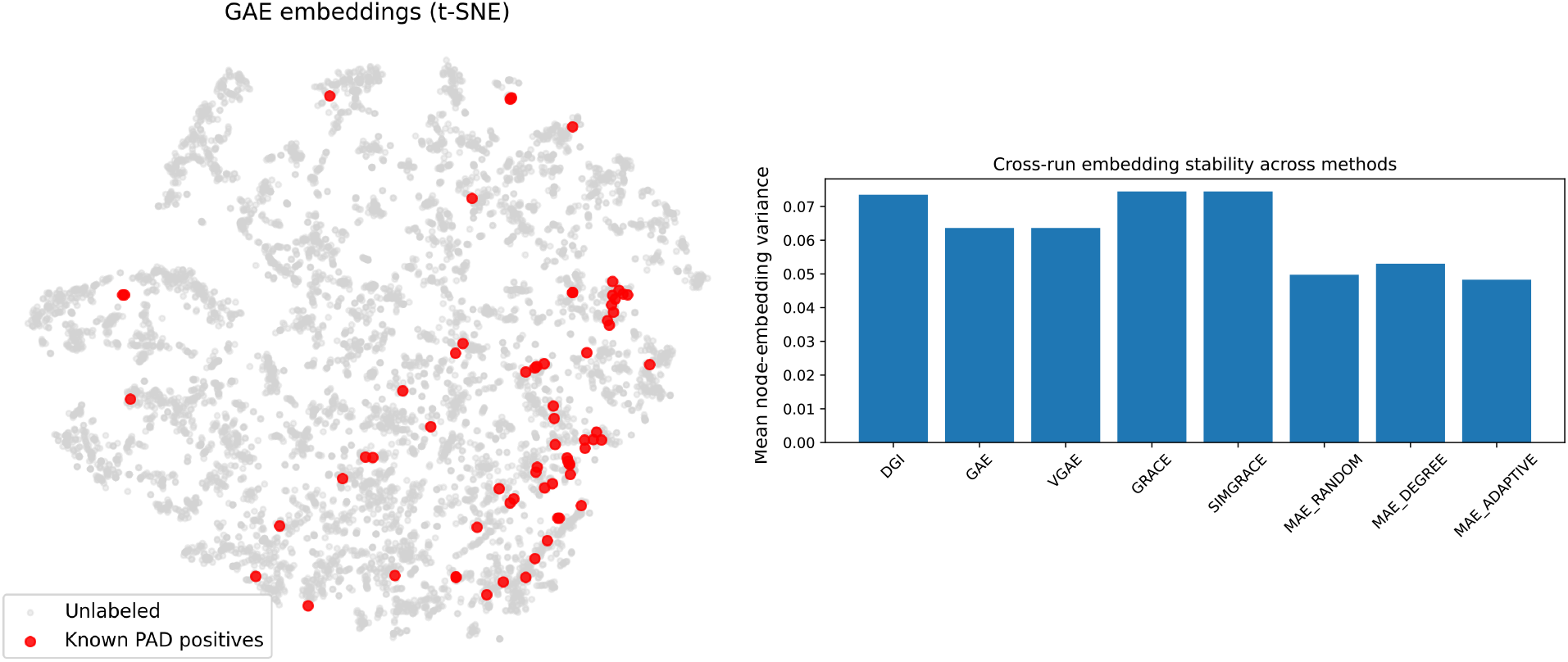
Embedding stability and t-SNE projection. (A) Cross-run embedding stability across all eight unsupervised methods, quantified as mean node-wise standard deviation across random initialisations. Lower values indicate greater reproducibility. (B) Representative two-dimensional t-SNE projection of GAE node embeddings with candidate stratification. Known PAD-associated proteins are shown in red, probability-heavy candidates in blue, and novelty-heavy candidates in green.

**Table S1.**
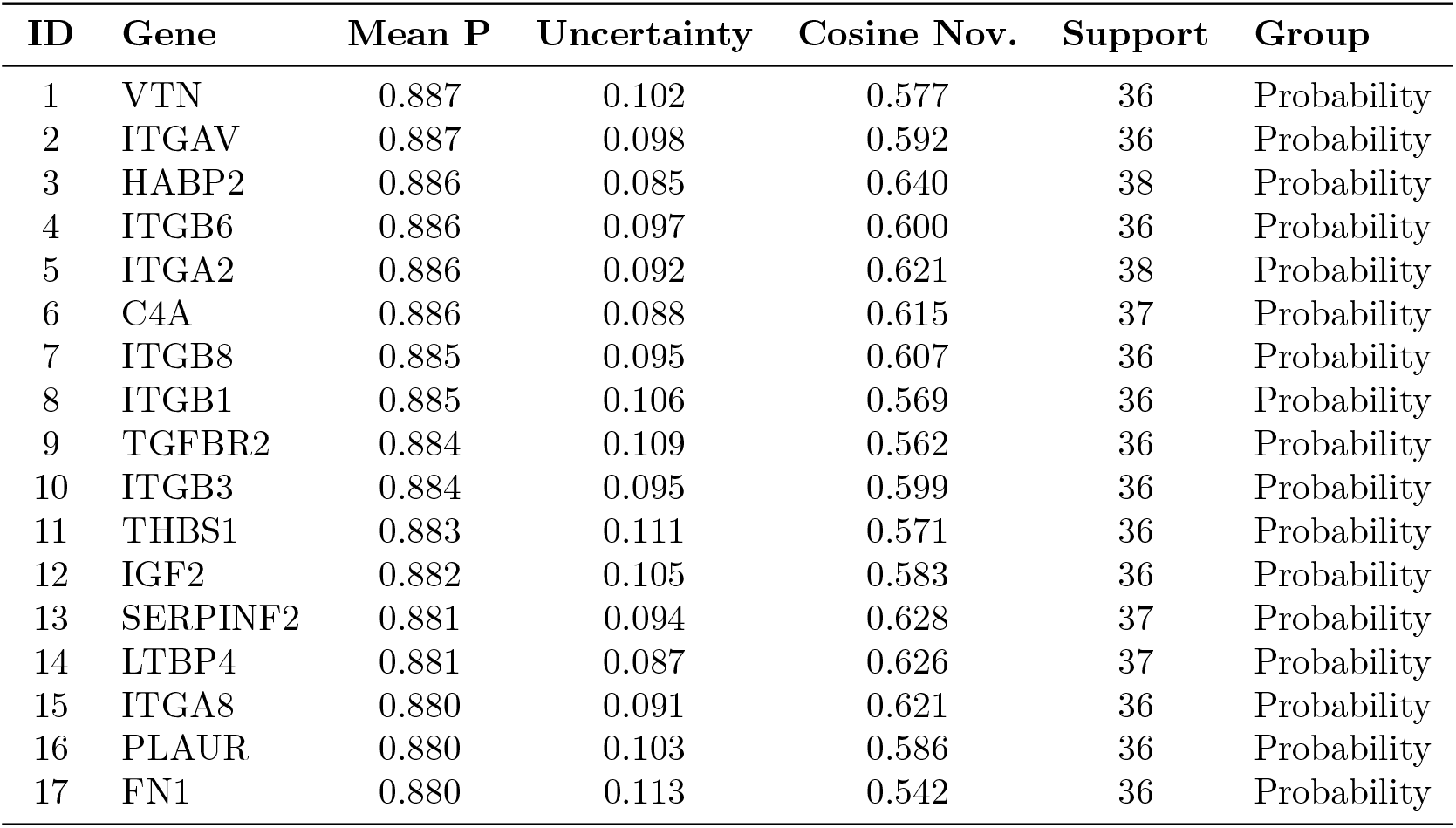
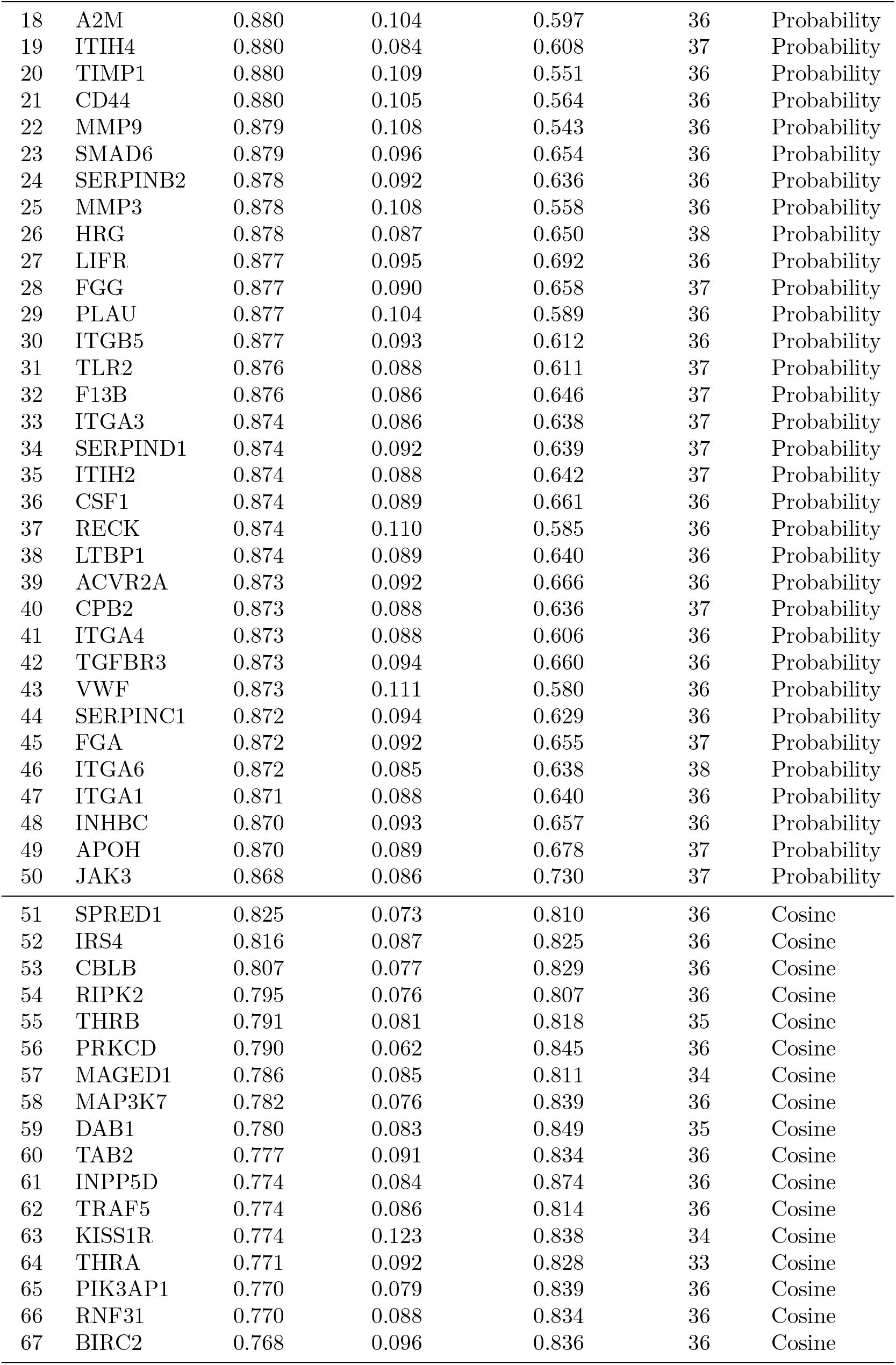
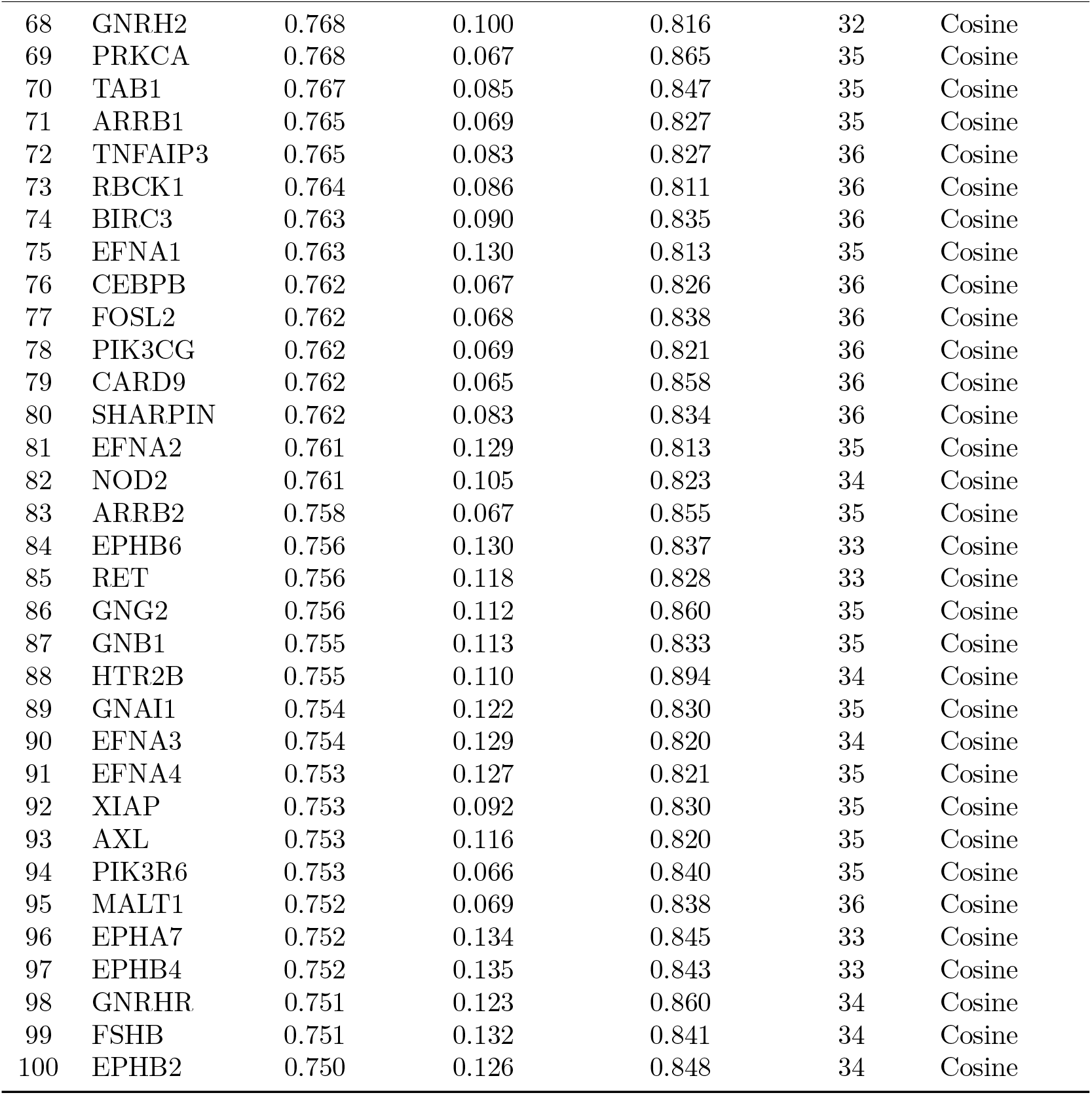
Full list of prioritised novel PAD biomarker candidates. All 100 candidates with ensemble probability, epistemic uncertainty, cosine novelty, classifier support count, and group assignment. IDs 1–50 are probability-heavy (P); IDs 51–100 are cosine-heavy (C).

| ID | Gene | Mean P | Uncertainty | Cosine Nov. | Support | Group |
| --- | --- | --- | --- | --- | --- | --- |
| 1 | VTN | 0.887 | 0.102 | 0.577 | 36 | Probability |
| 2 | ITGAV | 0.887 | 0.098 | 0.592 | 36 | Probability |
| 3 | HABP2 | 0.886 | 0.085 | 0.640 | 38 | Probability |
| 4 | ITGB6 | 0.886 | 0.097 | 0.600 | 36 | Probability |
| 5 | ITGA2 | 0.886 | 0.092 | 0.621 | 38 | Probability |
| 6 | C4A | 0.886 | 0.088 | 0.615 | 37 | Probability |
| 7 | ITGB8 | 0.885 | 0.095 | 0.607 | 36 | Probability |
| 8 | ITGB1 | 0.885 | 0.106 | 0.569 | 36 | Probability |
| 9 | TGFBR2 | 0.884 | 0.109 | 0.562 | 36 | Probability |
| 10 | ITGB3 | 0.884 | 0.095 | 0.599 | 36 | Probability |
| 11 | THBS1 | 0.883 | 0.111 | 0.571 | 36 | Probability |
| 12 | IGF2 | 0.882 | 0.105 | 0.583 | 36 | Probability |
| 13 | SERPINF2 | 0.881 | 0.094 | 0.628 | 37 | Probability |
| 14 | LTBP4 | 0.881 | 0.087 | 0.626 | 37 | Probability |
| 15 | ITGA8 | 0.880 | 0.091 | 0.621 | 36 | Probability |
| 16 | PLAUR | 0.880 | 0.103 | 0.586 | 36 | Probability |
| 17 | FN1 | 0.880 | 0.113 | 0.542 | 36 | Probability |
| 18 | A2M | 0.880 | 0.104 | 0.597 | 36 | Probability |
| 19 | ITIH4 | 0.880 | 0.084 | 0.608 | 37 | Probability |
| 20 | TIMP1 | 0.880 | 0.109 | 0.551 | 36 | Probability |
| 21 | CD44 | 0.880 | 0.105 | 0.564 | 36 | Probability |
| 22 | MMP9 | 0.879 | 0.108 | 0.543 | 36 | Probability |
| 23 | SMAD6 | 0.879 | 0.096 | 0.654 | 36 | Probability |
| 24 | SERPINB2 | 0.878 | 0.092 | 0.636 | 36 | Probability |
| 25 | MMP3 | 0.878 | 0.108 | 0.558 | 36 | Probability |
| 26 | HRG | 0.878 | 0.087 | 0.650 | 38 | Probability |
| 27 | LIFR | 0.877 | 0.095 | 0.692 | 36 | Probability |
| 28 | FGG | 0.877 | 0.090 | 0.658 | 37 | Probability |
| 29 | PLAU | 0.877 | 0.104 | 0.589 | 36 | Probability |
| 30 | ITGB5 | 0.877 | 0.093 | 0.612 | 36 | Probability |
| 31 | TLR2 | 0.876 | 0.088 | 0.611 | 37 | Probability |
| 32 | F13B | 0.876 | 0.086 | 0.646 | 37 | Probability |
| 33 | ITGA3 | 0.874 | 0.086 | 0.638 | 37 | Probability |
| 34 | SERPIND1 | 0.874 | 0.092 | 0.639 | 37 | Probability |
| 35 | ITIH2 | 0.874 | 0.088 | 0.642 | 37 | Probability |
| 36 | CSF1 | 0.874 | 0.089 | 0.661 | 36 | Probability |
| 37 | RECK | 0.874 | 0.110 | 0.585 | 36 | Probability |
| 38 | LTBP1 | 0.874 | 0.089 | 0.640 | 36 | Probability |
| 39 | ACVR2A | 0.873 | 0.092 | 0.666 | 36 | Probability |
| 40 | CPB2 | 0.873 | 0.088 | 0.636 | 37 | Probability |
| 41 | ITGA4 | 0.873 | 0.088 | 0.606 | 36 | Probability |
| 42 | TGFBR3 | 0.873 | 0.094 | 0.660 | 36 | Probability |
| 43 | VWF | 0.873 | 0.111 | 0.580 | 36 | Probability |
| 44 | SERPINC1 | 0.872 | 0.094 | 0.629 | 36 | Probability |
| 45 | FGA | 0.872 | 0.092 | 0.655 | 37 | Probability |
| 46 | ITGA6 | 0.872 | 0.085 | 0.638 | 38 | Probability |
| 47 | ITGA1 | 0.871 | 0.088 | 0.640 | 36 | Probability |
| 48 | INHBC | 0.870 | 0.093 | 0.657 | 36 | Probability |
| 49 | APOH | 0.870 | 0.089 | 0.678 | 37 | Probability |
| 50 | JAK3 | 0.868 | 0.086 | 0.730 | 37 | Probability |
| 51 | SPRED1 | 0.825 | 0.073 | 0.810 | 36 | Cosine |
| 52 | IRS4 | 0.816 | 0.087 | 0.825 | 36 | Cosine |
| 53 | CBLB | 0.807 | 0.077 | 0.829 | 36 | Cosine |
| 54 | RIPK2 | 0.795 | 0.076 | 0.807 | 36 | Cosine |
| 55 | THRB | 0.791 | 0.081 | 0.818 | 35 | Cosine |
| 56 | PRKCD | 0.790 | 0.062 | 0.845 | 36 | Cosine |
| 57 | MAGED1 | 0.786 | 0.085 | 0.811 | 34 | Cosine |
| 58 | MAP3K7 | 0.782 | 0.076 | 0.839 | 36 | Cosine |
| 59 | DAB1 | 0.780 | 0.083 | 0.849 | 35 | Cosine |
| 60 | TAB2 | 0.777 | 0.091 | 0.834 | 36 | Cosine |
| 61 | INPP5D | 0.774 | 0.084 | 0.874 | 36 | Cosine |
| 62 | TRAF5 | 0.774 | 0.086 | 0.814 | 36 | Cosine |
| 63 | KISS1R | 0.774 | 0.123 | 0.838 | 34 | Cosine |
| 64 | THRA | 0.771 | 0.092 | 0.828 | 33 | Cosine |
| 65 | PIK3AP1 | 0.770 | 0.079 | 0.839 | 36 | Cosine |
| 66 | RNF31 | 0.770 | 0.088 | 0.834 | 36 | Cosine |
| 67 | BIRC2 | 0.768 | 0.096 | 0.836 | 36 | Cosine |
| 68 | GNRH2 | 0.768 | 0.100 | 0.816 | 32 | Cosine |
| 69 | PRKCA | 0.768 | 0.067 | 0.865 | 35 | Cosine |
| 70 | TAB1 | 0.767 | 0.085 | 0.847 | 35 | Cosine |
| 71 | ARRB1 | 0.765 | 0.069 | 0.827 | 35 | Cosine |
| 72 | TNFAIP3 | 0.765 | 0.083 | 0.827 | 36 | Cosine |
| 73 | RBCK1 | 0.764 | 0.086 | 0.811 | 36 | Cosine |
| 74 | BIRC3 | 0.763 | 0.090 | 0.835 | 36 | Cosine |
| 75 | EFNA1 | 0.763 | 0.130 | 0.813 | 35 | Cosine |
| 76 | CEBPB | 0.762 | 0.067 | 0.826 | 36 | Cosine |
| 77 | FOSL2 | 0.762 | 0.068 | 0.838 | 36 | Cosine |
| 78 | PIK3CG | 0.762 | 0.069 | 0.821 | 36 | Cosine |
| 79 | CARD9 | 0.762 | 0.065 | 0.858 | 36 | Cosine |
| 80 | SHARPIN | 0.762 | 0.083 | 0.834 | 36 | Cosine |
| 81 | EFNA2 | 0.761 | 0.129 | 0.813 | 35 | Cosine |
| 82 | NOD2 | 0.761 | 0.105 | 0.823 | 34 | Cosine |
| 83 | ARRB2 | 0.758 | 0.067 | 0.855 | 35 | Cosine |
| 84 | EPHB6 | 0.756 | 0.130 | 0.837 | 33 | Cosine |
| 85 | RET | 0.756 | 0.118 | 0.828 | 33 | Cosine |
| 86 | GNG2 | 0.756 | 0.112 | 0.860 | 35 | Cosine |
| 87 | GNB1 | 0.755 | 0.113 | 0.833 | 35 | Cosine |
| 88 | HTR2B | 0.755 | 0.110 | 0.894 | 34 | Cosine |
| 89 | GNAI1 | 0.754 | 0.122 | 0.830 | 35 | Cosine |
| 90 | EFNA3 | 0.754 | 0.129 | 0.820 | 34 | Cosine |
| 91 | EFNA4 | 0.753 | 0.127 | 0.821 | 35 | Cosine |
| 92 | XIAP | 0.753 | 0.092 | 0.830 | 35 | Cosine |
| 93 | AXL | 0.753 | 0.116 | 0.820 | 35 | Cosine |
| 94 | PIK3R6 | 0.753 | 0.066 | 0.840 | 35 | Cosine |
| 95 | MALT1 | 0.752 | 0.069 | 0.838 | 36 | Cosine |
| 96 | EPHA7 | 0.752 | 0.134 | 0.845 | 33 | Cosine |
| 97 | EPHB4 | 0.752 | 0.135 | 0.843 | 33 | Cosine |
| 98 | GNRHR | 0.751 | 0.123 | 0.860 | 34 | Cosine |
| 99 | FSHB | 0.751 | 0.132 | 0.841 | 34 | Cosine |
| 100 | EPHB2 | 0.750 | 0.126 | 0.848 | 34 | Cosine |

### S1 Text: Supplementary Methods

#### S1.1 GATv2 encoder architecture

We trained a two-layer Graph Attention Network v2 (GATv2) [28] as the shared encoder for all unsupervised models. Both node features **X** ∈ ℝ^*N* ×*d*^ and edge features **e**_*ij*_ ∈ ℝ^*k*^ are incorporated into the message-passing operation:

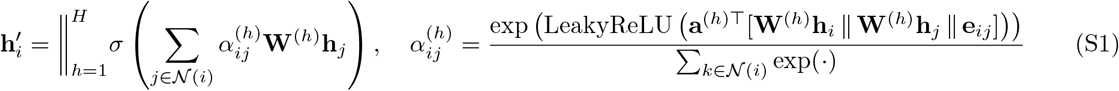

where *H* is the number of attention heads and ∥ denotes concatenation. The encoder uses *H* = 4 attention heads, hidden dimension 256, and outputs 128-dimensional embeddings per node. Dropout of 0.1 is applied during training. The Adam optimizer is used with learning rate 10^−3^ and weight decay 5 × 10^−4^.

**Fig S2.**
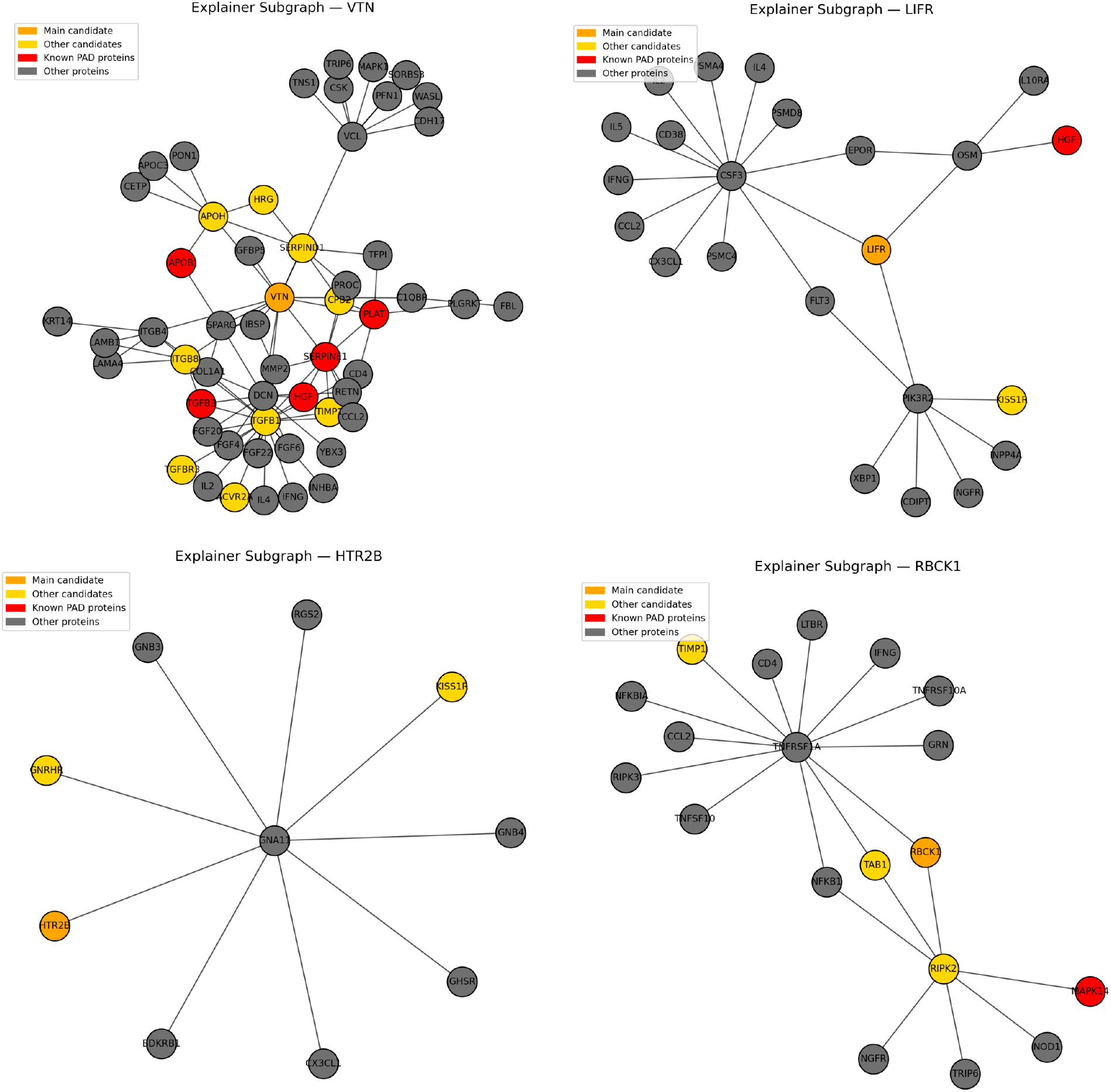
Representative GNNExplainer subgraphs. Shown are two probability-heavy candidates (VTN, LIFR) and two novelty-heavy candidates (HTR2B, RBCK1). Node colors indicate candidate proteins (orange), other candidates (yellow), known PAD-associated proteins (red), and background proteins (gray). While some high-confidence candidates are embedded within PAD-enriched neighbourhoods (e.g., VTN), others achieve strong support through structurally influential but PAD-sparse contexts (e.g., LIFR). Novelty-heavy candidates similarly span PAD-distant network configurations, illustrating the diversity of structural motifs underlying confident and exploratory predictions.

#### S1.2 Unsupervised training objectives

##### Deep Graph Infomax (DGI)

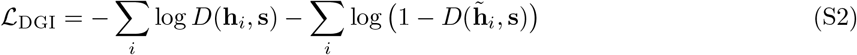

where 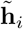 are embeddings from a corrupted graph and *D* is a bilinear discriminator function.

**Fig S3.**
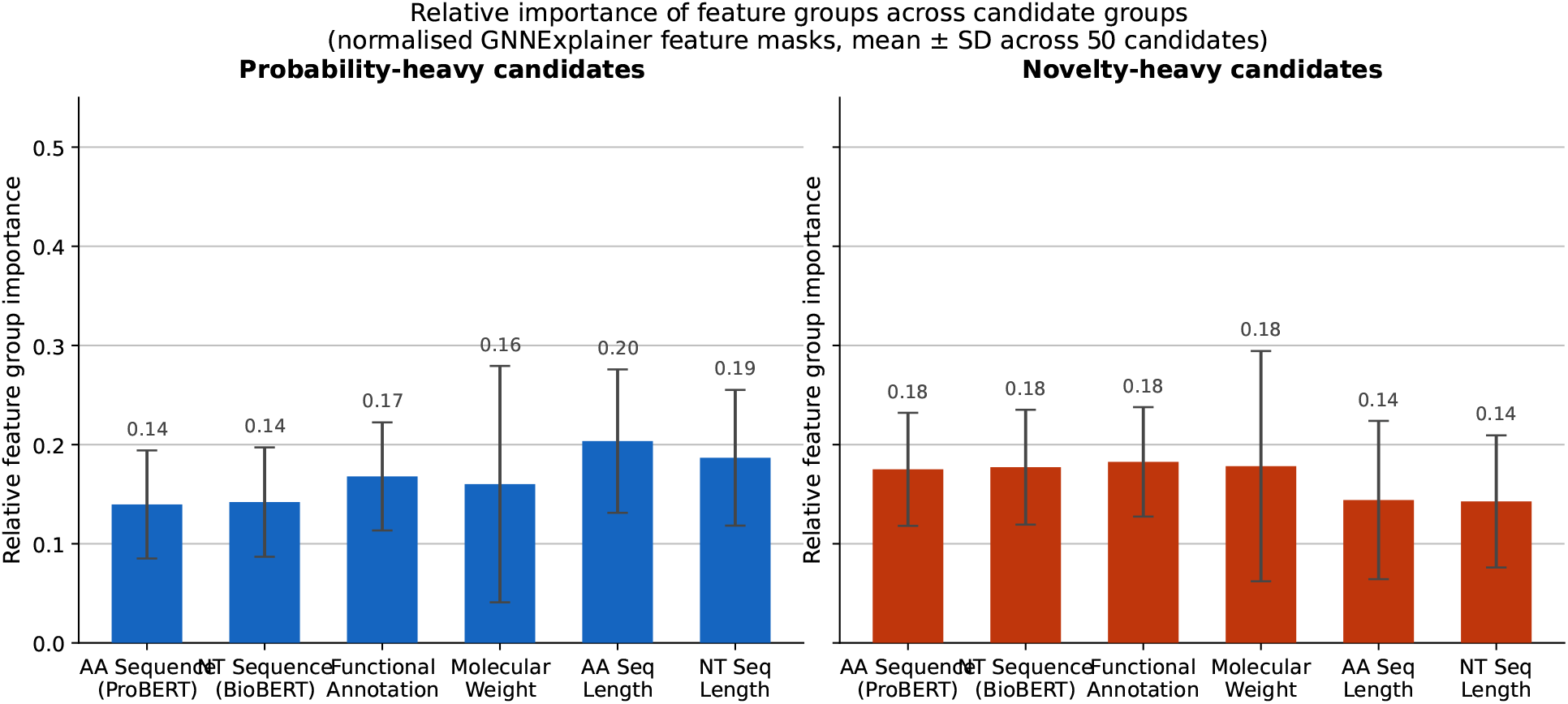
Relative importance of feature groups across candidate groups. Normalised GNNExplainer feature mask importance aggregated by feature group for probability-heavy (blue) and novelty-heavy (red) candidates. Values represent mean ± SD across 50 candidates per group. Feature groups correspond to the six multimodal protein representations: amino acid sequence embeddings (ProBERT), nucleotide sequence embeddings (BioBERT), functional annotation embeddings, molecular weight, amino acid sequence length, and nucleotide sequence length.

**Table S2.**
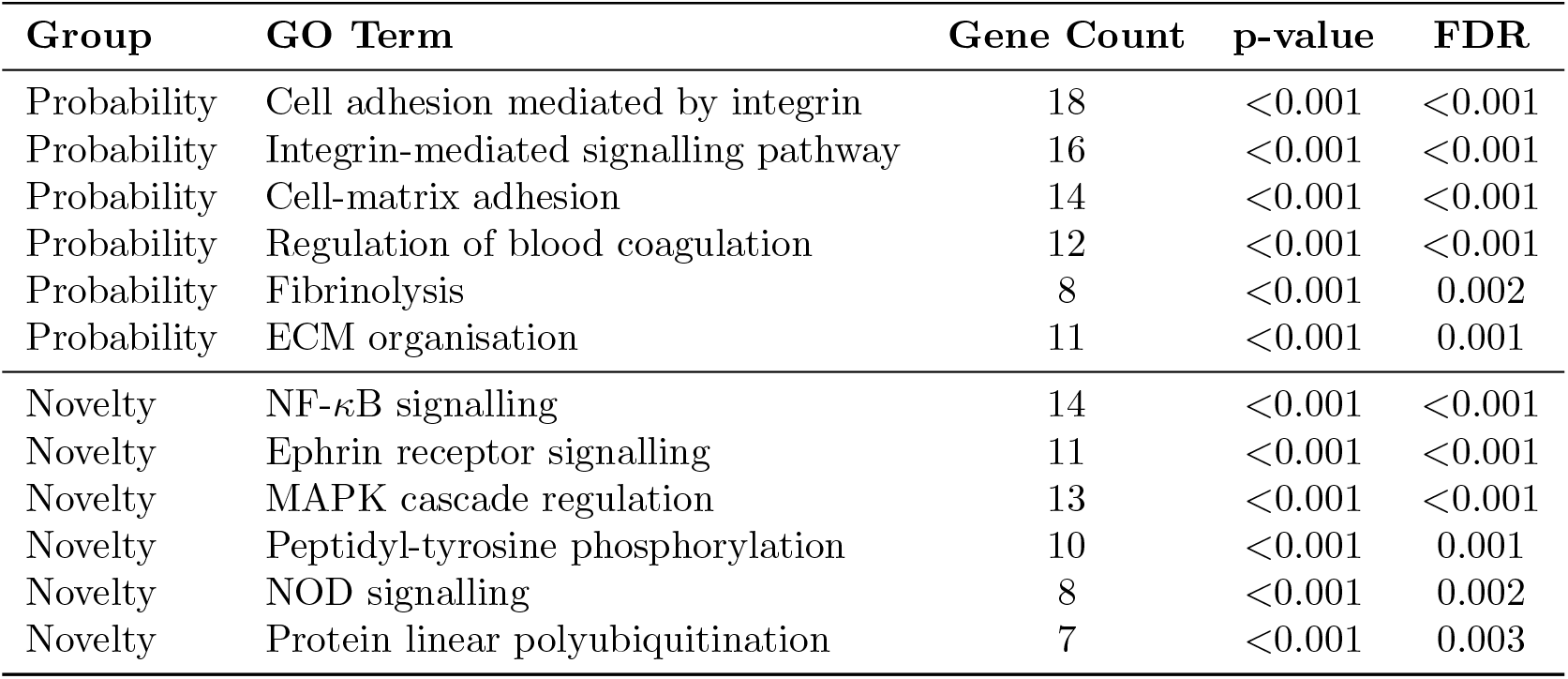
Pathway enrichment results. Complete Gene Ontology biological process enrichment for probability-heavy and novelty-heavy candidate groups. [To be populated with enrichment output — GO term, gene count, p-value, FDR, gene list per term.]

| Group | GO Term | Gene Count | p-value | FDR |
| --- | --- | --- | --- | --- |
| Probability | Cell adhesion mediated by integrin | 18 | <0.001 | <0.001 |
| Probability | Integrin-mediated signalling pathway | 16 | <0.001 | <0.001 |
| Probability | Cell-matrix adhesion | 14 | <0.001 | <0.001 |
| Probability | Regulation of blood coagulation | 12 | <0.001 | <0.001 |
| Probability | Fibrinolysis | 8 | <0.001 | 0.002 |
| Probability | ECM organisation | 11 | <0.001 | 0.001 |
| Novelty | NF-κB signalling | 14 | <0.001 | <0.001 |
| Novelty | Ephrin receptor signalling | 11 | <0.001 | <0.001 |
| Novelty | MAPK cascade regulation | 13 | <0.001 | <0.001 |
| Novelty | Peptidyl-tyrosine phosphorylation | 10 | <0.001 | 0.001 |
| Novelty | NOD signalling | 8 | <0.001 | 0.002 |
| Novelty | Protein linear polyubiquitination | 7 | <0.001 | 0.003 |

##### Graph Autoencoder (GAE) and Variational GAE (VGAE)

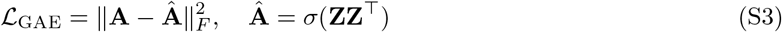

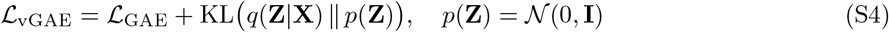

**Table S3.**
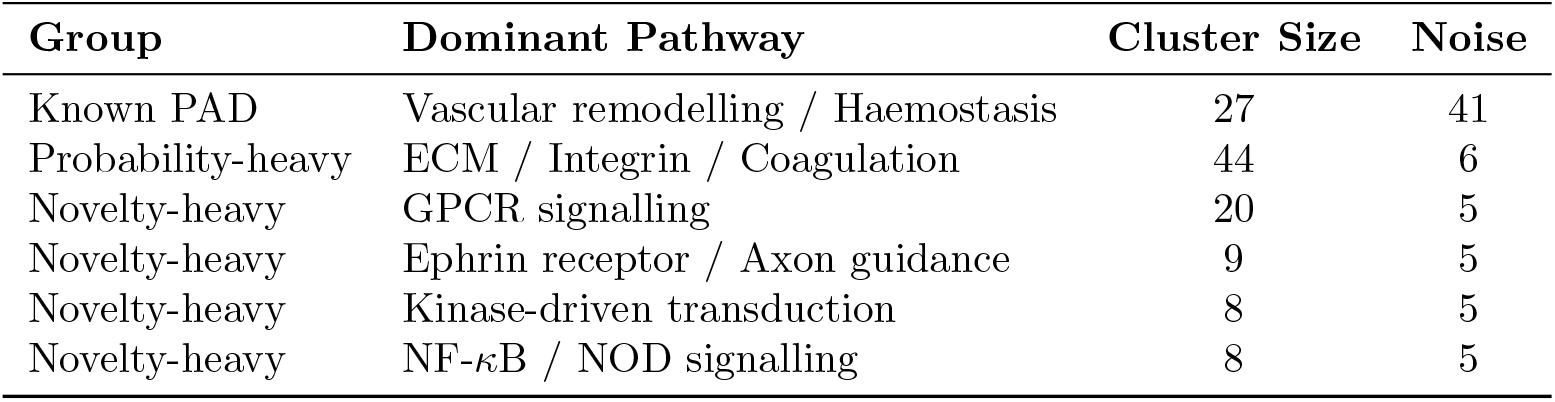
Clustering analysis. Cluster assignments and dominant pathway annotations for known PAD proteins, probability-heavy candidates, and novelty-heavy candidates.

| Group | Dominant Pathway | Cluster Size | Noise |
| --- | --- | --- | --- |
| Known PAD | Vascular remodelling / Haemostasis | 27 | 41 |
| Probability-heavy | ECM / Integrin / Coagulation | 44 | 6 |
| Novelty-heavy | GPCR signalling | 20 | 5 |
| Novelty-heavy | Ephrin receptor / Axon guidance | 9 | 5 |
| Novelty-heavy | Kinase-driven transduction | 8 | 5 |
| Novelty-heavy | NF-κB / NOD signalling | 8 | 5 |

##### GRACE and SimGRACE

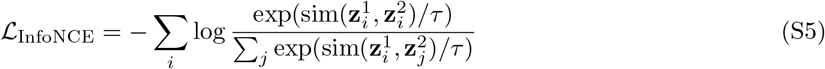

where sim is cosine similarity and *τ* = 0.5.

##### Masked Autoencoder (MAE)

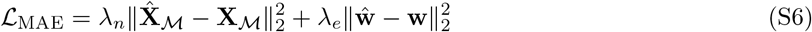

where *λ*_*n*_ = 1.0 and *λ*_*e*_ = 0.5. Masking ratio is 0.3 across all three strategies (random, degree-based, adaptive).

#### S1.3 Reliable negative identification

##### Heuristic methods

**Spy method**. A fraction *π* = 0.15 of known positives is injected into 𝒰 as spies. A Gaussian Naïve Bayes classifier is trained to discriminate spies from unlabelled nodes. All unlabelled nodes with posterior probability lower than the maximum posterior of any spy are retained as reliable negative candidates.

*k***-means distance filter**. Nodes are clustered via *k*-means with *k* = 20. Unlabelled nodes in non-positive clusters are candidate RNs, ranked by mean Euclidean distance to all known positives.

*k***-NN distance ranking**. Mean distance to *k* = 5 nearest known positives is computed for each unlabelled node.

**Generative PU model**. A single-component Gaussian mixture model is fitted to positive embeddings; unlabelled nodes with lowest log-likelihood are selected.

**CCRNE**. Unlabelled nodes from *k*-means clusters containing no positives are selected.

##### Uncertainty-based RN refinement

Each candidate RN node *i* is scored by three metrics:

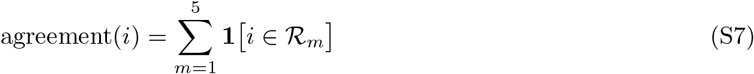

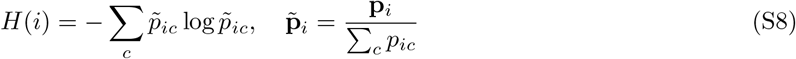

with 1 − *H*(*i*) as confidence, and mean Euclidean distance to *k* = 5 nearest positives. Each score is min-max normalised and the final confidence is their unweighted mean. The top |𝒫 | = 68 candidates are retained.

#### S1.4 Iterative PU classifier training

For MC Dropout and BNN, *S* = 30 stochastic forward passes are drawn:

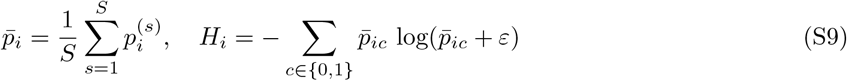

New RNs are added as: 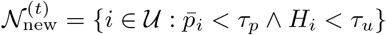 with *τ*_*p*_ = 0.2 and *τ*_*u*_ = 0.05. For classical models, nodes with *p*_*i*_ *< τ*_*p*_ are added (*τ*_*p*_ = 0.1 for RF).

Stopping criteria: (a) fewer than 5 new RNs, (b) Δ*F*_1_ *<* 10^−3^, or (c) false negative rate on known positives exceeds 5%. Maximum iterations: 10.

#### S1.5 Final uncertainty quantification

For MC Dropout and BNN:

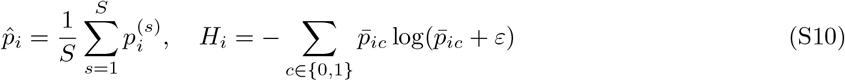

For deterministic classifiers:

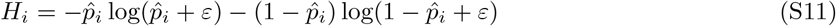

where *ε* = 10^−8^.

#### S1.6 GNNExplainer configuration

Configuration: 200 optimisation epochs, learning rate 0.01, edge and feature mask regularisation 0.01, top-*k* edge retention *k* = 10, *k*-hop neighbourhood = 2.

#### S1.7 Explanation evaluation metrics

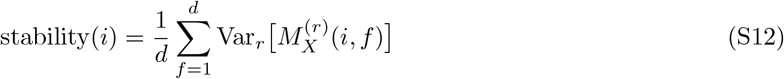

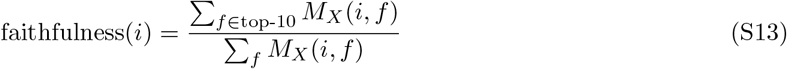

#### S1.8 Hyperparameter summary

**Table S4.**
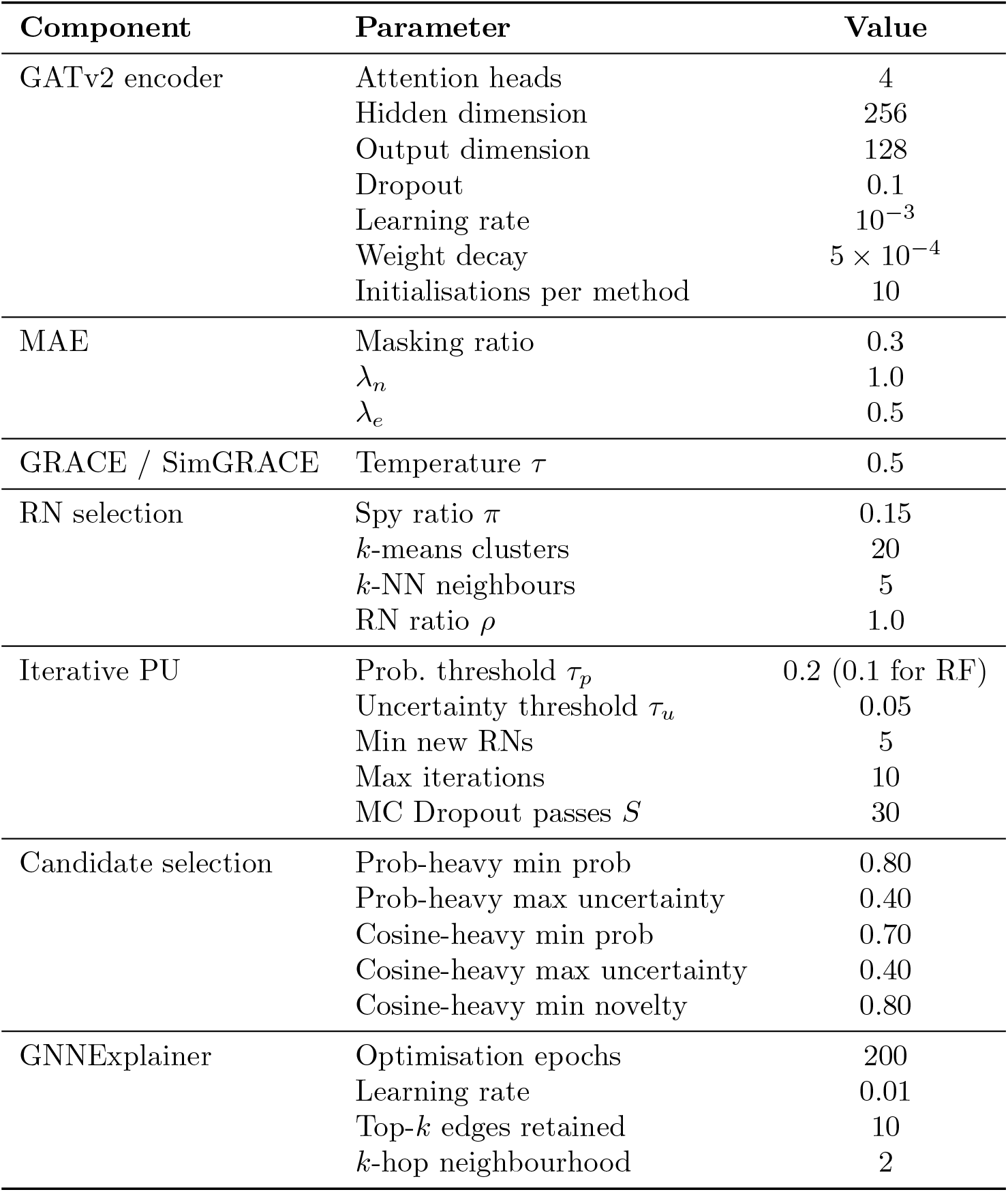
Hyperparameter settings for all model components.

## References

1. Song P, Rudan D, Zhu Y, Fowkes FJ, Rahimi K, Fowkes FGR, et al. Global, regional, and national prevalence and risk factors for peripheral artery disease in 2015: an updated systematic review and analysis. The Lancet Global Health. 2019;7(8):e1020–30.

2. Fowkes FGR, Aboyans V, Fowkes FJ, McDermott MM, Sampson UK, Criqui MH. Peripheral artery disease: epidemiology and global perspectives. Nature Reviews Cardiology. 2017;14(3):156–70.

3. McDermott MM, Liu K, Greenland P, Guralnik JM, Criqui MH, Chan C, et al. Functional decline in peripheral arterial disease: associations with the ankle brachial index and leg symptoms. Jama. 2004;292(4):453–61.

4. Newman AB, Shemanski L, Manolio TA, Cushman M, Mittelmark M, Polak JF, et al. Ankle-arm index as a predictor of cardiovascular disease and mortality in the Cardiovascular Health Study. Arteriosclerosis, thrombosis, and vascular biology. 1999;19(3):538–45.

5. Criqui MH, Langer RD, Fronek A, Feigelson HS, Klauber MR, McCann TJ, et al. Mortality over a period of 10 years in patients with peripheral arterial disease. New England Journal of Medicine. 1992;326(6):381–6.

6. Gornik HL, Creager MA. Contemporary management of peripheral arterial disease: I. Cardiovascular risk-factor modification. Cleveland Clinic journal of medicine. 2006;73:S30–7.

7. Hirsch AT, Haskal ZJ, Hertzer NR, Bakal CW, Creager MA, Halperin JL, et al. ACC/AHA 2005 practice guidelines for the management of patients with peripheral arterial disease (lower extremity, renal, mesenteric, and abdominal aortic) a collaborative report from the American Association for Vascular Surgery/Society for Vascular Surgery,* Society for Cardiovascular Angiography and Interventions, Society for Vascular Medicine and Biology, Society of Interventional Radiology, and the ACC/AHA Task Force on Practice Guidelines (writing committee to develop guidelines for the management of patients with peripheral arterial disease): endorsed by the American Association of Cardiovascular and Pulmonary Rehabilitation; National Heart, Lung, and Blood Institute; Society for Vascular Nursing; TransAtlantic Inter-Society Consensus; and Vascular Disease Foundation. circulation. 2006;113(11):e463–654.

8. McDermott MM, Greenland P, Liu K, Guralnik JM, Criqui MH, Dolan NC, et al. Leg symptoms in peripheral arterial disease: associated clinical characteristics and functional impairment. Jama. 2001;286(13):1599–606.

9. Norgren L, Hiatt WR, Dormandy JA, Nehler MR, Harris KA, Fowkes FGR, et al. Inter-society consensus for the management of peripheral arterial disease (TASC II). Journal of vascular surgery. 2007;45(1):S5–S67.

10. Cooke JP, Wilson AM. Biomarkers of peripheral arterial disease. Journal of the American College of Cardiology. 2010;55(19):2017–23.

11. Khan H, Girdharry NR, Massin SZ, Abu-Raisi M, Saposnik G, Mamdani M, et al. Current Prognostic Biomarkers for Peripheral Arterial Disease: A Comprehensive Systematic Review of the Literature. Metabolites. 2025;15(4):224.

12. Fung ET, Wilson AM, Zhang F, Harris N, Edwards KA, Olin JW, et al. A biomarker panel for peripheral arterial disease. Vascular Medicine. 2008;13(3):217–24.

13. Saenz-Pipaon G, Martinez-Aguilar E, Orbe J, Gonzalez Miqueo A, Fernandez-Alonso L, Paramo JA, et al. The role of circulating biomarkers in peripheral arterial disease. International Journal of Molecular Sciences. 2021;22(7):3601.

14. Ferrucci L, Candia J, Ubaida-Mohien C, Lyashkov A, Banskota N, Leeuwenburgh C, et al. Transcriptomic and proteomic of gastrocnemius muscle in peripheral artery disease. Circulation research. 2023;132(11):1428–43.

15. Jain I, Oropeza BP, Huang NF. Multiomics analyses of peripheral artery disease muscle biopsies. Lippincott Williams & Wilkins Hagerstown, MD; 2023.

16. Hasin Y, Seldin M, Lusis A. Multi-omics approaches to disease. Genome biology. 2017;18(1):83.

17. Eraslan G, Avsec Ž, Gagneur J, Theis FJ. Deep learning: new computational modelling techniques for genomics. Nature reviews genetics. 2019;20(7):389–403.

18. Zitnik M, Agrawal M, Leskovec J. Modeling polypharmacy side effects with graph convolutional networks. Bioinformatics. 2018;34(13):i457–66.

19. Cowen L, Ideker T, Raphael BJ, Sharan R. Network propagation: a universal amplifier of genetic associations. Nature Reviews Genetics. 2017;18(9):551–62.

20. Yang P, Li XL, Mei JP, Kwoh CK, Ng SK. Positive-unlabeled learning for disease gene identification. Bioinformatics. 2012;28(20):2640–7.

21. Bekker J, Davis J. Learning from positive and unlabeled data: A survey. Machine learning. 2020;109(4):719–60.

22. Yang P, Li X, Chua HN, Kwoh CK, Ng SK. Ensemble positive unlabeled learning for disease gene identification. PloS one. 2014;9(5):e97079.

23. Mordelet F, Vert JP. Prodige: Prioritization of disease genes with multitask machine learning from positive and unlabeled examples. BMC bioinformatics. 2011;12(1):389.

24. Stolfi P, Mastropietro A, Pasculli G, Tieri P, Vergni D. NIAPU: network-informed adaptive positive-unlabeled learning for disease gene identification. Bioinformatics. 2023;39(2):btac848.

25. Szklarczyk D, Kirsch R, Koutrouli M, Nastou K, Mehryary F, Hachilif R, et al. The STRING database in 2023: protein–protein association networks and functional enrichment analyses for any sequenced genome of interest. Nucleic acids research. 2023;51(D1):D638–46.

26. Elnaggar A, Heinzinger M, Dallago C, Rehawi G, Wang Y, Jones L, et al. ProtTrans: Towards Cracking the Language of Life’s Code Through Self-Supervised Deep Learning and High Performance Computing. bioRxiv. 2020. Available from: https://www.biorxiv.org/content/early/2020/07/21/2020.07.12.199554. arXiv:https://www.biorxiv.org/content/early/2020/07/21/2020.07.12.199554.full.pdf. doi:10.1101/2020.07.12.199554.

27. Lee J, Yoon W, Kim S, Kim D, Kim S, So CH, et al. BioBERT: a pre-trained biomedical language representation model for biomedical text mining. Bioinformatics. 2020;36(4):1234–40.

28. Brody S, Alon U, Yahav E. How attentive are graph attention networks? arXiv preprint arXiv:210514491. 2021.

29. Veličković P, Cucurull G, Casanova A, Romero A, Lio P, Bengio Y. Graph attention networks. arXiv preprint arXiv:171010903. 2017.

30. Veličković P, Fedus W, Hamilton WL, Lio P, Bengio Y, Hjelm RD. Deep graph infomax. arXiv preprint arXiv:180910341. 2018.

31. Kipf TN, Welling M. Variational graph auto-encoders. arXiv preprint arXiv:161107308. 2016.

32. Zhu Y, Xu Y, Yu F, Liu Q, Wu S, Wang L. Deep graph contrastive representation learning. arXiv preprint arXiv:200604131. 2020.

33. Xia J, Wu L, Chen J, Hu B, Li SZ. Simgrace: A simple framework for graph contrastive learning without data augmentation. In: Proceedings of the ACM web conference 2022; 2022. p. 1070–9.

34. He K, Chen X, Xie S, Li Y, Dollár P, Girshick R. Masked autoencoders are scalable vision learners. In: Proceedings of the IEEE/CVF conference on computer vision and pattern recognition; 2022. p. 16000–9.

35. Gal Y, Ghahramani Z. Dropout as a bayesian approximation: Representing model uncertainty in deep learning. In: international conference on machine learning. PMLR; 2016. p. 1050–9.

36. Lakshminarayanan B, Pritzel A, Blundell C. Simple and scalable predictive uncertainty estimation using deep ensembles. Advances in neural information processing systems. 2017;30.

37. Ying Z, Bourgeois D, You J, Zitnik M, Leskovec J. Gnnexplainer: Generating explanations for graph neural networks. Advances in neural information processing systems. 2019;32.

38. Agarwal C, Zitnik M, Lakkaraju H. Probing gnn explainers: A rigorous theoretical and empirical analysis of gnn explanation methods. In: International conference on artificial intelligence and statistics. PMLR; 2022. p. 8969–96.

39. Gillis J, Pavlidis P. The impact of multifunctional genes on” guilt by association” analysis. PloS one. 2011;6(2):e17258.

40. Hüllermeier E, Waegeman W. Aleatoric and epistemic uncertainty in machine learning: An introduction to concepts and methods. Machine learning. 2021;110(3):457–506.

41. Abdar M, Khosravi A, Islam SMS, Acharya UR, Vasilakos AV. The need for quantification of uncertainty in artificial intelligence for clinical data analysis: increasing the level of trust in the decision-making process. IEEE Systems, Man, and Cybernetics Magazine. 2022;8(3):28–40.

42. Begoli E, Bhattacharya T, Kusnezov D. The need for uncertainty quantification in machine-assisted medical decision making. Nature Machine Intelligence. 2019;1(1):20–3.

43. Sendak MP, D’Arcy J, Kashyap S, Gao M, Nichols M, Corey K, et al. A path for translation of machine learning products into healthcare delivery. EMJ Innov. 2020;10:19–00172.

44. Krychtiuk K, Kastl S, Speidl W, Wojta J. Inflammation and coagulation in atherosclerosis. Hämostaseologie. 2013;33(04):269–82.

45. De Martin R, Hoeth M, Hofer-Warbinek R, Schmid JA. The transcription factor NF-*κ*B and the regulation of vascular cell function. Arteriosclerosis, thrombosis, and vascular biology. 2000;20(11):e83–8.

46. Xu F, Ma J, Wang X, Wang X, Fang W, Sun J, et al. The role of G protein-coupled estrogen receptor (GPER) in vascular pathology and physiology. Biomolecules. 2023;13(9):1410.

47. Zheng S, Sun F, Tian X, Zhu Z, Wang Y, Zheng W, et al. Roles of Eph/ephrin signaling pathway in repair and regeneration for ischemic cerebrovascular and cardiovascular diseases. Journal of Neurorestoratology. 2023;11(1):100040.

48. Fu S, Zhao H, Shi J, Abzhanov A, Crawford K, Ohno-Machado L, et al. Peripheral arterial occlusive disease: global gene expression analyses suggest a major role for immune and inflammatory responses. Bmc Genomics. 2008;9(1):369.

49. Li T, Li X, Feng Y, Dong G, Wang Y, Yang J. The role of matrix metalloproteinase-9 in atherosclerotic plaque instability. Mediators of inflammation. 2020;2020(1):3872367.

50. Zhang K, Li M, Yin L, Fu G, Liu Z. Role of thrombospondin-1 and thrombospondin-2 in cardiovascular diseases. International journal of molecular medicine. 2020;45(5):1275–93.

51. Habib A, Petrucci G, Rocca B. Pathophysiology of thrombosis in peripheral artery disease. Current Vascular Pharmacology. 2020;18(3):204–14.

52. Van Der Bom J, Bots M, Haverkate F, Meijer P, Hofman A, Kluft C, et al. Activation products of the haemostatic system in coronary, cerebrovascular and peripheral arterial disease. Thrombosis and haemostasis. 2001;85(02):234–9.

